# Tumor Genotype Dictates Mitochondrial and Immune Vulnerabilities in Liver Cancer

**DOI:** 10.1101/2025.09.10.675369

**Authors:** Gokhan Unlu, Alon Millet, Khando Wangdu, Romain Donné, Ranya Erdal, Nicole L. DelGaudio, Beste Uygur, Vyom Shah, Kevin Cho, Antonia Fecke, Feyza Cansiz, Zeynep Cagla Tarcan, Michael Isay-Del Viscio, Ece Kilic, Isabel Kurth, Henrik Molina, Albert Sickmann, Olca Basturk, Gary Patti, Semir Beyaz, Karl W. Smith, Amaia Lujambio, Alpaslan Tasdogan, Sohail F. Tavazoie, Kıvanç Birsoy

**Affiliations:** Laboratory of Metabolic Regulation and Genetics, The Rockefeller University, New York, NY 10065, USA; Laboratory of Systems Cancer Biology, The Rockefeller University, New York, NY, 10065, USA; Tri-Institutional Program in Computational Biology and Medicine, The Rockefeller University, New York, NY 10065, USA; Department of Oncological Sciences, Liver Cancer Program, Division of Liver Diseases, Department of Medicine, Tisch Cancer Institute, Icahn School of Medicine at Mount Sinai, New York, NY, 10029, USA; Medical Scientist Training Program, Hacettepe University Faculty of Medicine, Ankara 06230, Turkey; Cold Spring Harbor Laboratory, Cold Spring Harbor, New York, USA; Department of Chemistry, Washington University, St Louis, MO, USA; Leibniz-Institut für Analytische Wissenschaften - ISAS - e.V., Dortmund, Germany; Department of Dermatology, University Hospital Essen & German Cancer Consortium (DKTK), Partner Site, Essen, Germany; Department of Pathology and Laboratory Medicine Memorial Sloan Kettering Cancer Center, New York, New York, USA; The Proteomics Resource Center, The Rockefeller University, New York, NY, 10065, USA; Inspirna, Inc., 30-02 48^th^ Ave., Long Island City, NY 11101, USA

## Abstract

Although oncogenic alterations influence tumor metabolism, how they impose distinct metabolic programs within a shared tissue context remains poorly defined. Here, we developed a rapid mitochondrial profiling platform to compare metabolites and proteins in genetic models of primary liver cancer (PLC). Analyses of six genetically distinct PLCs revealed that mitochondrial energy metabolism is largely dictated by oncogene identity. *Kras*-driven tumors required creatine metabolism to buffer energy demands during early tumorigenesis, whereas *c-MYC*-driven tumors relied on oxidative phosphorylation. Among *c-MYC*-driven PLCs, *Pten*-deficient tumors accumulated mitochondrial phosphoethanolamine, a precursor for phosphatidylethanolamine (PE) synthesis. Inhibition of PE synthesis selectively impaired the growth of *Pten*-deficient tumors and extended survival, in part through enhanced infiltration of CD8⁺ T cells and sensitization to TNFα-mediated cytotoxicity. Mechanistically, loss of PE elevated surface TNF receptor 2 (TNFR2), promoting TNFα signaling and pro-inflammatory response. These findings uncover genotype-specific mitochondrial metabolic liabilities and establish PE synthesis as a tumor-intrinsic mechanism of immune evasion in PLC.

## INTRODUCTION

To support their growth and proliferation, cancer cells rely on metabolic pathways that supply energy, generate biosynthetic precursors, and maintain redox balance^1^. Many of these pathways operate downstream of oncogenic signaling cascades frequently dysregulated in cancers ^2–4^. For example, *PI3K* stimulates glycolysis and lipid synthesis to support anabolic growth, whereas *MYC* directly transactivates lactate dehydrogenase A (LDHA), linking oncogenic signaling to aerobic glycolysis for rapid ATP generation ^5, 6^. Beyond oncogenes, tumor suppressors such as *p53*, when mutated, also rewire metabolic pathways by altering glycolytic flux and antioxidant pathways to support cellular fitness^7^. Although oncogenic drivers can activate overlapping metabolic programs, their outputs vary depending on cell type and tissue of origin. Indeed, comparative analyses of human tumors have shown that metabolic signatures often reflect those of the corresponding normal tissue, suggesting that malignant transformation does not fully erase lineage-specific metabolic features ^8^. This is exemplified by lung adenocarcinomas, which exhibit greater dependence on branched-chain amino acid (BCAA) metabolism than pancreatic cancers, despite harboring similar oncogenic lesions ^9^. Because both genotype and tissue context influence tumor metabolism, isolating the metabolic effects of specific oncogenic alterations requires a controlled, tissue-matched system. This complexity is particularly evident in primary liver cancer (PLC), a malignancy with high mortality and limited treatment options. PLCs exhibit broad genetic heterogeneity ^10^, with recurrent alterations in oncogenes and tumor suppressors such as c-*MYC*, *KRAS*, *PTEN*, *KEAP1* and *TP53,* yet how these mutations influence cancer metabolism *in vivo* remains unclear ^11, 12^.

## RESULTS

### Integrative mitochondrial profiling reveals genotype-specific metabolic programs

To study how distinct oncogenic alterations influence cancer cell metabolism *in vivo*, we developed a rapid and cell type–specific method to isolate mitochondria from genetically engineered mouse models (GEMMs) that recapitulate tumorigenesis from normal liver tissue. Given their central role in coordinating metabolic inputs and dynamically exchanging metabolites with the cytosol, mitochondria serve as a robust surrogate for assessing cellular metabolic state^13^. To enable selective isolation from cancer cells, our approach adapts a previously developed mitochondrially localized HA epitope (mito-tag) for immunopurification (mito-IP)^14^. We engineered vectors to bicistronically express an oncogene and a luciferase-fused mito-tag using an internal ribosomal entry site (IRES), allowing for simultaneous tumor induction, selective expression in cancer cells and non-invasive tumor imaging (Figures 1a and S1a,b). These constructs were delivered into C57BL/6J mice via hydrodynamic tail vein injection (HDTVi) to induce liver tumors^15, 16^. We confirmed cancer cell–specific expression and mitochondrial localization of the mito-tag by immunofluorescence (Figure S1c-f), and validated the robustness of mitochondrial purification for proteomic and metabolomic analyses (Figure S1g,h). Building on our platform, we next sought to model six genetically defined tumor types commonly observed in human PLCs using oncogene expression vectors delivered via HDTVi ^15–18^. These vectors co-expressed c-*MYC* with sgRNAs targeting *Trp53*, *Pten*, *Keap1*, or *Kmt2c* to induce loss-of-function mutations; c-*MYC* in combination with an N-terminally truncated, constitutively active β-catenin ^15^ (*Ctnnb1,* referred to as B-cat Δ90) modeling hepatocellular carcinoma (HCC); or Kras^G12D^ with *Trp53* sgRNAs to model intrahepatic cholangiocarcinoma (ICC)^19^. Mitochondria from CMV-Cre; mito-tag^fl/fl^ mouse livers ^20^ served as healthy controls for comparative analyses (Figure 1b).

**Figure 1.**
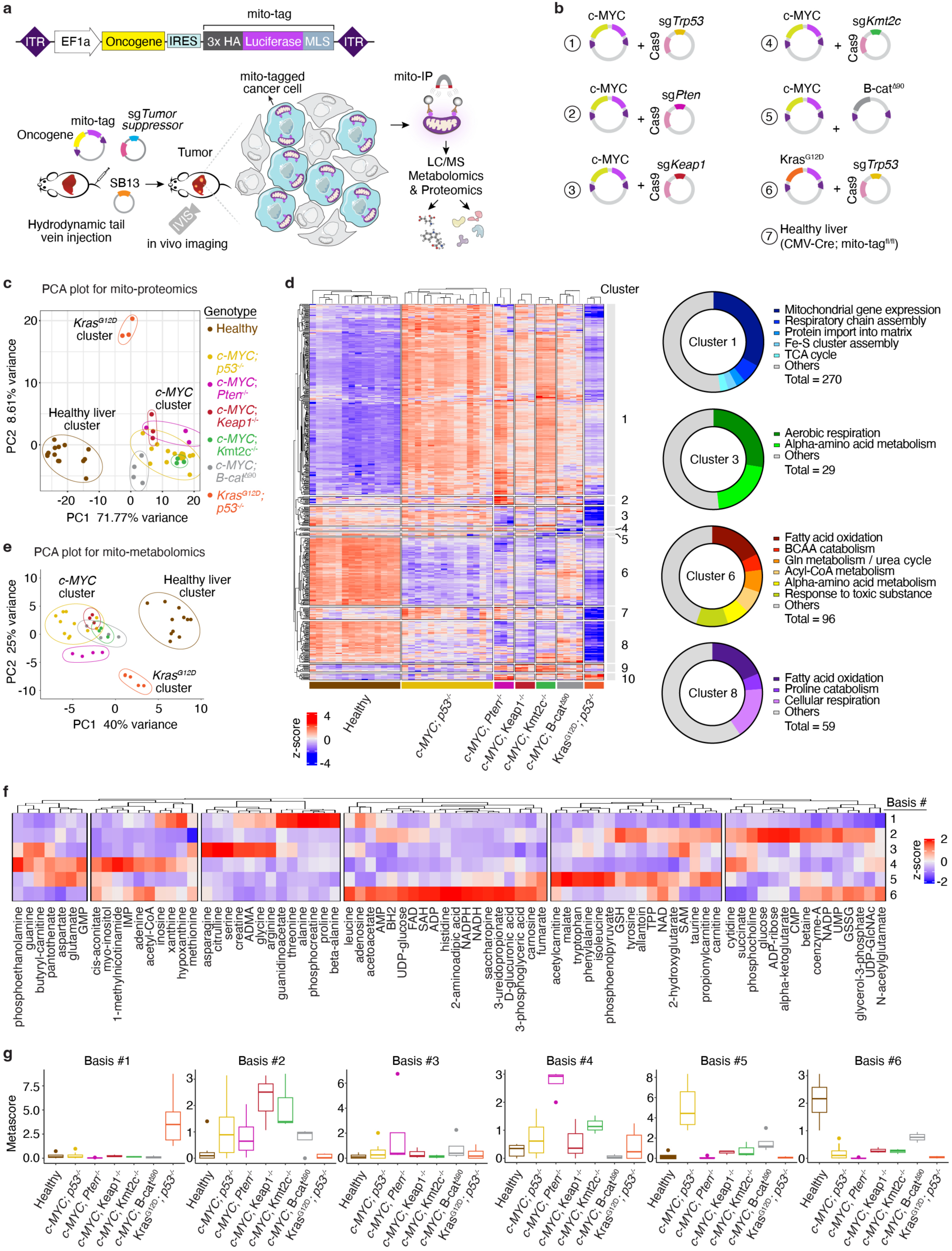
Rapid mitochondrial immunopurification method to profile cancer cell mitochondria from tumors. a. Overview of cell-type specific labeling of liver cancer cells *in vivo* and rapid mitochondrial immunocapture method from tumors. Transposon-based genomic integration construct for bicistronic expression of oncogene and mito-tag is depicted on top. The versatile genetic toolkit enables tumor induction, *in vivo* monitoring of tumor growth by bioluminescence and molecular profiling by metabolomics and proteomics. IVIS: *in vivo* imaging system. LC/MS: liquid chromatography / mass spectroscopy. ITR: inverted terminal repeat, IRES: internal ribosomal entry site, MLS: mitochondrial localization signal from OMP25, SB13: sleeping beauty transposase 13. b. Genetic constructs for modeling PLC with common mutations by HDTVi. c. PCA plot for mitochondrial proteome of healthy liver and genetic PLC models. d. Heatmap showing median normalized changes in mitochondrial proteins isolated from healthy livers and different tumor models, as indicated below by colored bars. Rows and columns are clustered by hierarchical clustering. Cluster labels are indicated by numbers on the right. e. PCA plot for mitochondrial metabolome of healthy liver and genetic PLC models f. Heatmap of mitochondrial metabolites categorized into 6 non-negative matrix factorization (NMF) basis matrices. Z-scores are indicated across blue-to-red color scale. g. Box whisker plots of metascores calculated for each tumor type and NMF basis matrix.

To define mitochondrial proteomic programs across these genotypes, we performed proteomic analysis on immunopurified mitochondria from tumors and healthy liver, detecting 859 proteins (513 annotated as mitochondrial; Figure S2a). Principal component analysis (PCA) revealed three distinct clusters: (1) healthy liver, (2) *Kras^G12D^*-driven tumors, and (3) *c-MYC*-driven tumors (Figure 1c), indicating that oncogene identity is a dominant determinant of mitochondrial protein composition. Hierarchical clustering further defined ten proteomic clusters with distinct expression patterns (Figure 1d and Table S1). Among the ten clusters, Cluster 1, enriched in *c-MYC* tumors, contained proteins involved in mitochondrial translation, the TCA cycle, and respiratory chain assembly for oxidative phosphorylation (OXPHOS) (Figures 1d and S2b,c). Cluster 3, containing aerobic respiration and alpha-amino acid metabolism components, was selectively downregulated in *Kras^G12D^*-driven tumors. Liver-enriched enzymes supporting normal hepatic functions such as fatty acid oxidation, BCAA catabolism, and the urea cycle (Clusters 6 and 8) were broadly suppressed across tumor types (Figure S2b), in line with prior work^10, 21^. Notably, sub-clustering within c*-MYC* tumors revealed additional stratification based on tumor suppressor status (Figure S2d), suggesting that tumor suppressor loss further modulates mitochondrial composition, albeit more subtly than oncogenic drivers. As a proof of principle, we tested whether these proteomic alterations represent functional dependencies in *c-MYC*–driven tumors, which exhibit selective enrichment of mitochondrial translation and OXPHOS proteins compared to *Kras^G12D^*-driven tumors and healthy liver (Figures 1d and S2b,c). Pharmacological inhibition of complex I using IACS-10759 markedly suppressed *c-MYC; p53^-/-^* tumor growth, while *Kras^G12D^; p53^-/-^* tumors were unaffected (Figure S2e). Similarly, genetic depletion of *Mrps22*, a mitochondrial ribosomal subunit elevated in *c-MYC* tumors, impaired tumor growth (Figure S2f), indicating a dependency on mitochondrial translation. These findings identify mitochondrial translation and OXPHOS as genotype-specific metabolic vulnerabilities in *c-MYC*– driven PLC.

We next profiled the mitochondrial metabolome from the same models to get a clearer view of deregulated metabolic pathways using LC/MS-based analysis of polar mitochondrial metabolites (Figure S3a). PCA again revealed distinct clustering of *c-MYC*- and *Kras*-driven tumors, with healthy liver forming a separate group (Figures 1e and S3b). Non-negative matrix factorization method (NMF) provides a means to investigate the complex mitochondrial metabolomics dataset in an interpretable and simplified manner in form of basis groups; therefore, we generated six optimal NMF bases that captured the entire dataset, spanning all genotypes (Figure 1f,g). Basis #1, enriched in *Kras^G12D^; p53^-/-^* tumors, included phosphocreatine, guanidinoacetate, proline, and arginine (Figures 1f,g and S3c,d). Basis #2, elevated across all *c-MYC* tumors, included TCA intermediates (e.g., α-ketoglutarate, succinate) in line with the mitochondrial proteomics profiling (Figure 1d, cluster 1), and glutathione, with GSH levels highest in *Keap1*-null tumors, consistent with KEAP1 as a negative regulator of NRF2, a transcriptional activator of GSH synthesis and antioxidant pathways^22^ (Figures 1f,g and S3c,d). In contrast, Basis #6 reflected healthy liver metabolism, including lysine degradation intermediates saccharopine and 2-aminoadipate (Figures 1f,g and S3c). These findings reveal complementary proteomic and metabolomic programs that distinguish PLCs by genotype and suggest potential metabolic liabilities.

### Creatine metabolism is a selective metabolic liability for *Kras^G12D^*-driven PLCs

To gain a deeper insight into genotype-specific metabolic alterations, we developed an integrative approach that cross-references mitochondrial metabolomics and proteomics datasets. Specifically, we selected the top enriched proteins and metabolites in each tumor model and filtered metabolite-protein pairs that participate in the same metabolic pathway. Correlation and interaction scores were then computed based on enrichment within each tumor genotype (Figure S4a and Table S2). Among these, creatine metabolism emerged as the top enriched pathway in *Kras^G12D^; p53^-/-^* tumors (Figures 2a and S4a,b). Creatine is synthesized from arginine and glycine via GATM, followed by conversion of GAA to creatine^23^, which is phosphorylated by creatine kinase (CK) to form P-creatine (Figure 2b), a high-energy buffer for rapid ATP regeneration upon demand. Mitochondrial metabolic analysis consistently detected significant enrichment of creatine metabolism intermediates in *Kras^G12D^; p53^-/-^* tumors compared to both *c-MYC; p53^-/-^* tumors and healthy liver (Figures 2c and S3a,c,d). Supporting these findings, proteomics and immunoblotting assays revealed elevated levels of GATM in *Kras*-mutant tumor mitochondria and in a human KRAS-mutant ICC sample (Figures 2d and S4c,d). Spatial metabolomics using MALDI-MSI validated *in situ* enrichment of creatine and P-creatine, but not the vascular marker heme, in *Kras^G12D^; p53^-/-^* tumor regions (Figures 2e and S4e). Additionally, pantothenate exhibited selective accumulation in c*-MYC; p53^-/-^*tumors (Figure S4f), consistent with our mito-metabolomics data (Figure S3d) and a previous report ^24^. Finally, to confirm functional activation of creatine biosynthesis, we performed *in vivo* metabolite tracing using isotope labeled [^13^C_6_]-arginine and measured [^13^C_1_]-GAA as a readout of GATM function^25^, detecting increased [^13^C_1_]-GAA in tumors compared to adjacent healthy liver (Figure S5a).

**Figure 2.**
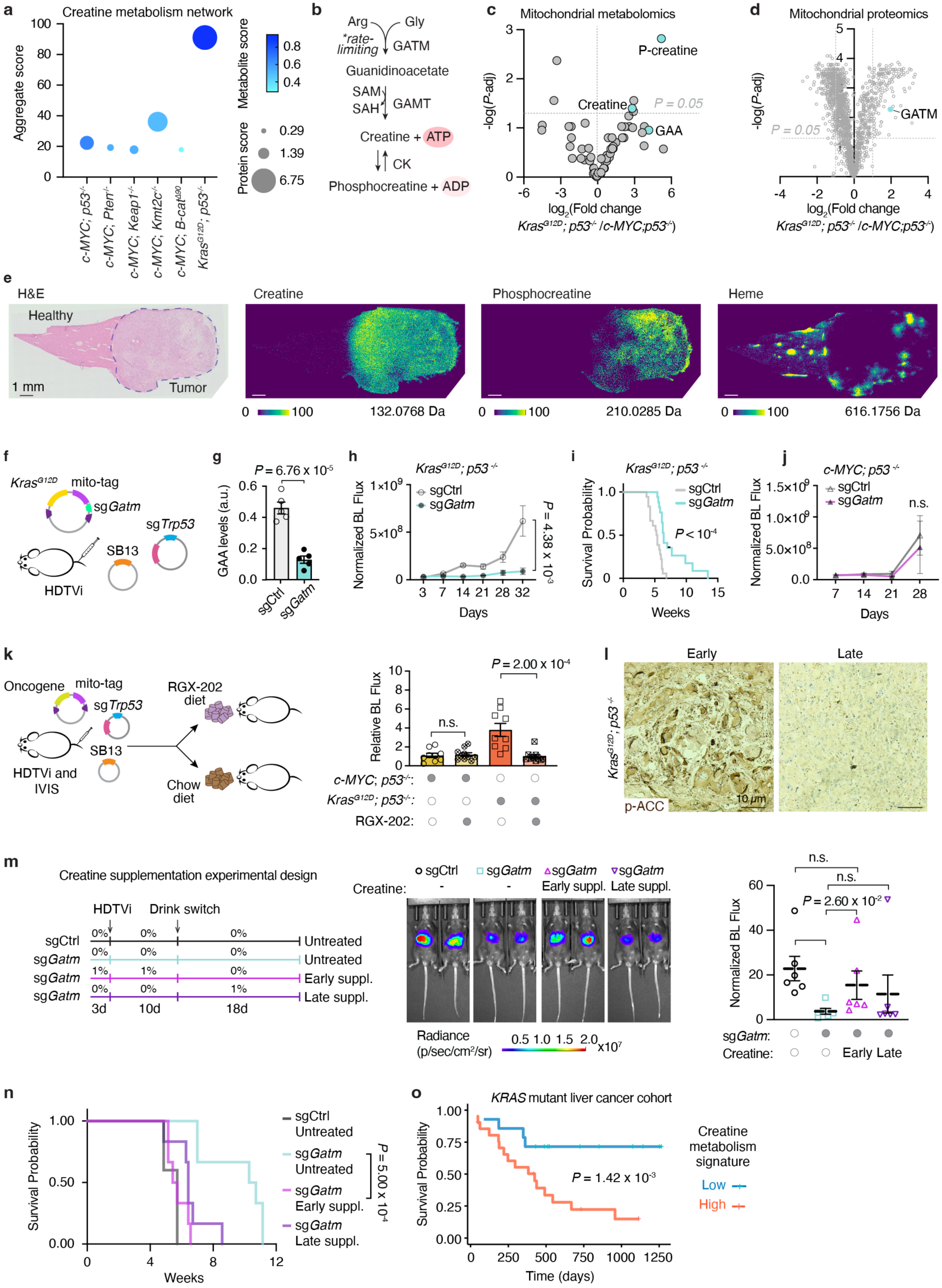
Creatine metabolism is a liability for *Kras^G12D^*-driven PLCs. a. Integrative metabolic network analysis results showing enrichment of creatine metabolism in *Kras^G12D^; p53^-/-^* tumors as compared to all other tumor models. b. Creatine biosynthesis pathway diagram. GATM is the rate limiting enzyme catalyzing the first and mitochondrial step of creatine biosynthesis. c. Volcano plot of mitochondrial metabolites showing comparison between *Kras^G12D^; p53^-/-^* and *c-MYC; p53^-/-^* tumors (log_2_-scale). -log *P*-value was calculated by Student’s t-test and false discovery rate (FDR)-adjusted by Benjamini-Hochberg. Confidence interval 95%. Horizontal dotted line demarcates significance at *P* = 0.05. GAA: guanidinoacetate. d. Volcano plot of mitochondrial proteins showing comparison between *Kras^G12D^; p53^-/-^* and *c-MYC; p53^-/-^* tumors (log_2_-scale). -log *P*-value was calculated by Student’s t-test and FDR-adjusted by Benjamini-Hochberg. Confidence interval 95%. Horizontal dotted line demarcates significance at p = 0.05. Vertical dotted lines represent boundaries for ≥ 2-fold change. e. H&E staining of mouse liver tissue with *Kras^G12D^; p53^-/-^* tumor. Tumor region is demarcated by dashed lines. MALDI-MSI images of creatine [M+H^+^], phosphocreatine [M+H^-^] and heme [M+H^+^] in *Kras^G12D^; p53^-/-^* tumor and healthy adjacent tissue, of the same specimen shown on the left H&E staining panel. Creatine and phosphocreatine signals show selective enrichment within tumor region, whereas heme signal intensity is higher around blood vessels across both tumor and adjacent tissues. Color bar below indicates % signal intensity. Depicted molecular ion mass is indicated below the image. Da: Dalton. Scale bar is 1 mm. f. HDTVi-based genetic depletion strategy for *Gatm* in PLC tumors. g. Mitochondrial GAA levels measured by LC/MS, in control (sgCtrl) and *Gatm*-depleted (sg*Gatm*) cancer cells isolated from tumors by cell type-specific mito-IP. *P*-value by Mann-Whitney U-test with 95% confidence interval. h. Normalized bioluminescence (BL) flux of *Kras^G12D^; p53^-/-^* tumors as measured by IVIS. Growth of control (sgCtrl) and *Gatm*-depleted (sg*Gatm*) tumors was compared by Mann-Whitney U-test with 95% confidence interval. i. Survival curve for control (sgCtrl) and *Gatm*-depleted (sg*Gatm*) *Kras^G12D^; p53^-/-^* tumor bearing mice. Curve comparison by Mantel-Cox test, confidence interval 95%. j. Normalized BL flux of *c-MYC; p53^-/-^* tumors as measured by IVIS. Growth of control (sgCtrl) and *Gatm*-depleted (sg*Gatm*) tumors was compared by Mann-Whitney U-test with 95% confidence interval. n.s.: not significant. k. Overview of RGX-202 diet experiment to test the growth of *Kras^G12D^; p53^-/-^* and *c-MYC; p53^-/-^* tumors. Bar plot showing relative BL flux of *c-MYC; p53^-/-^* and *Kras^G12D^; p53^-/-^* tumor growth. Curve comparison by Mann-Whitney U-test with 95% confidence interval. l. Immunohistochemistry analysis of P-ACC in early (10 days) and late (28 days) stage *Kras^G12D^; p53^-/-^* tumors. Scale bar is 10 *μ*m. m. Experimental design for creatine supplementation of mice bearing *Kras^G12D^; p53^-/-^* control (sgCtrl) or *Gatm*-depleted (sg*Gatm*) tumors. suppl.: supplementation. Middle panel, IVIS images of *Kras^G12D^; p53^-/-^* control (sgCtrl) or *Gatm*-depleted (sg*Gatm*) tumor bearing mice with or without creatine supplementation. On the right, plot displaying normalized BL flux (to 3-day timepoint) of *Kras^G12D^; p53^-/-^* tumors. Data were plotted as mean ± SEM; comparison by Mann-Whitney U-test with 95% confidence interval. n.s.: not significant. n. Survival curve for *Kras^G12D^; p53^-/-^* tumor bearing mice with or without creatine supplementation. Curve comparison by Mantel-Cox test, confidence interval 95%. o. Kaplan-Meier survival plot of ICC patients with *KRAS*-mutated tumors, based on composite transcript and protein expression scores of creatine metabolism genes. Curve comparison by Mantel-Cox test, confidence interval 95%.

Given its selective activation, we considered whether creatine biosynthesis is required for *Kras*-driven tumor growth. *Gatm* loss in *Kras^G12D^; p53^-/-^* tumors via HDTVi impaired tumor progression and extended survival (Figures 2f–I and S5b). In contrast, *c-MYC; p53^-/-^* tumors were unaffected by *Gatm* loss (Figure 2j). Creatine metabolism has recently been implicated in colon cancer metastasis, and a small molecule inhibitor of creatine metabolism, RGX-202 (ompenaclid), is under clinical development for advanced colorectal cancer^26, 27^. To test whether pharmacologic disruption of this pathway suppresses *Kras^G12D^*-driven PLCs, we fed *Kras^G12D^;p53^-/-^* and *c-MYC; p53^-/-^* tumor-bearing mice a modified diet containing RGX-202 (∼650 mg/kg, based on average daily consumption). *Kras^G12D^; p53^-/-^* tumors in mice fed RGX-202 progressed over three-fold more slowly than those in mice on control diets. In contrast, RGX-202 had no significant effect on *c-MYC*-driven tumor growth (Figure 2k). These findings suggest that oncogene-driven tumors exhibit distinct energy programs as metabolic liabilities: *c-MYC* tumors depend on OXPHOS (Figures 1d and S2c,e), whereas *Kras^G12D^*-driven tumors rely on creatine metabolism for energy buffering.

We next asked why creatine synthesis is selectively required in *Kras^G12D^*-driven tumors and considered a limitation in energy homeostasis. *Kras^G12D^*-driven tumors displayed a >50-fold increase in the AMP/ATP ratio compared to *c-MYC* driven tumors and normal liver (Figure S5c), indicating profound energy stress. Loss of *Gatm* further exacerbated this imbalance and activated AMP-activated protein kinase (AMPK), a central energy sensor that phosphorylates acetyl-CoA carboxylase (ACC) in response to energy stress. Indeed, phospho-ACC (P-ACC) levels were elevated in *Gatm*-deficient tumors (Figure S5d,e). Notably, phospho-ACC staining was strongest in early-stage tumors, indicating that energy stress is most pronounced during early tumor growth (Fig 2l). Consistently, RNA-seq analysis revealed enrichment of energy homeostasis pathways in early, but not late, tumors (Figure S5f), implicating creatine metabolism in buffering energy demand during early tumorigenesis. To directly test this early requirement for creatine, we supplemented mice bearing *Kras^G12D^; p53^-/-^* tumors with 1% creatine in drinking water either before (early) or after (late) tumor onset (Figure 2m). Metabolomic profiling confirmed elevated circulating creatine and its degradation product sarcosine in supplemented mice (Figure S6a). Strikingly, early, but not late, creatine supplementation restored *Gatm*-deficient tumor growth and reversed the associated survival benefit (Figure 2m,n), indicating a critical requirement for creatine during early tumor development. Notably, high expression of creatine metabolism genes was associated with significantly worse survival in a cohort of ICC patients with *KRAS*-mutant tumors ^28^ (Figures 2o and S6b). These findings identify creatine metabolism as a selective vulnerability that supports energy buffering and early tumor outgrowth in *Kras*-driven liver cancer.

### Essential role of PE synthesis in *PTEN*-null *c-MYC*-driven PLCs

To test whether tumor suppressors impose distinct metabolic liabilities, we next compared mitochondrial profiles across five *c-MYC*-driven tumor models. Among these, *c-MYC; Pten^-/-^* tumors exhibited a unique mitochondrial proteomic and metabolomic signature (Figures S2d and S3b), marked by elevated phosphoethanolamine (P-ethanolamine) (Figures 3a,b and S7a). P-ethanolamine is a key intermediate in the Kennedy pathway^29^, which drives the synthesis of phosphatidylethanolamine (PE), a major constituent of cellular membranes (Figure 3c). While P-ethanolamine can enter mitochondria and has been implicated in modulating respiration ^30–32^, its accumulation most likely reflects elevated cellular pools driven by enhanced PE biosynthesis in *Pten*-null tumors^33^. Indeed, lipidomic profiling confirmed a selective increase in several PE species in *c-MYC*; *Pten^-/-^* tumors (Figure S7b). We also assessed lipid droplet accumulation by Oil Red O staining and observed abundant lipid droplets in *c-MYC; Pten^-/-^* tumors (Figure 3d). Analysis of human PLCs revealed a similar pattern, with *PTEN*-mutant tumors displaying larger lipid droplets than PTEN wild-type cases (Figure 3e). These findings establish elevated PE synthesis and lipid storage as defining features of *Pten*-deficient tumors.

**Figure 3.**
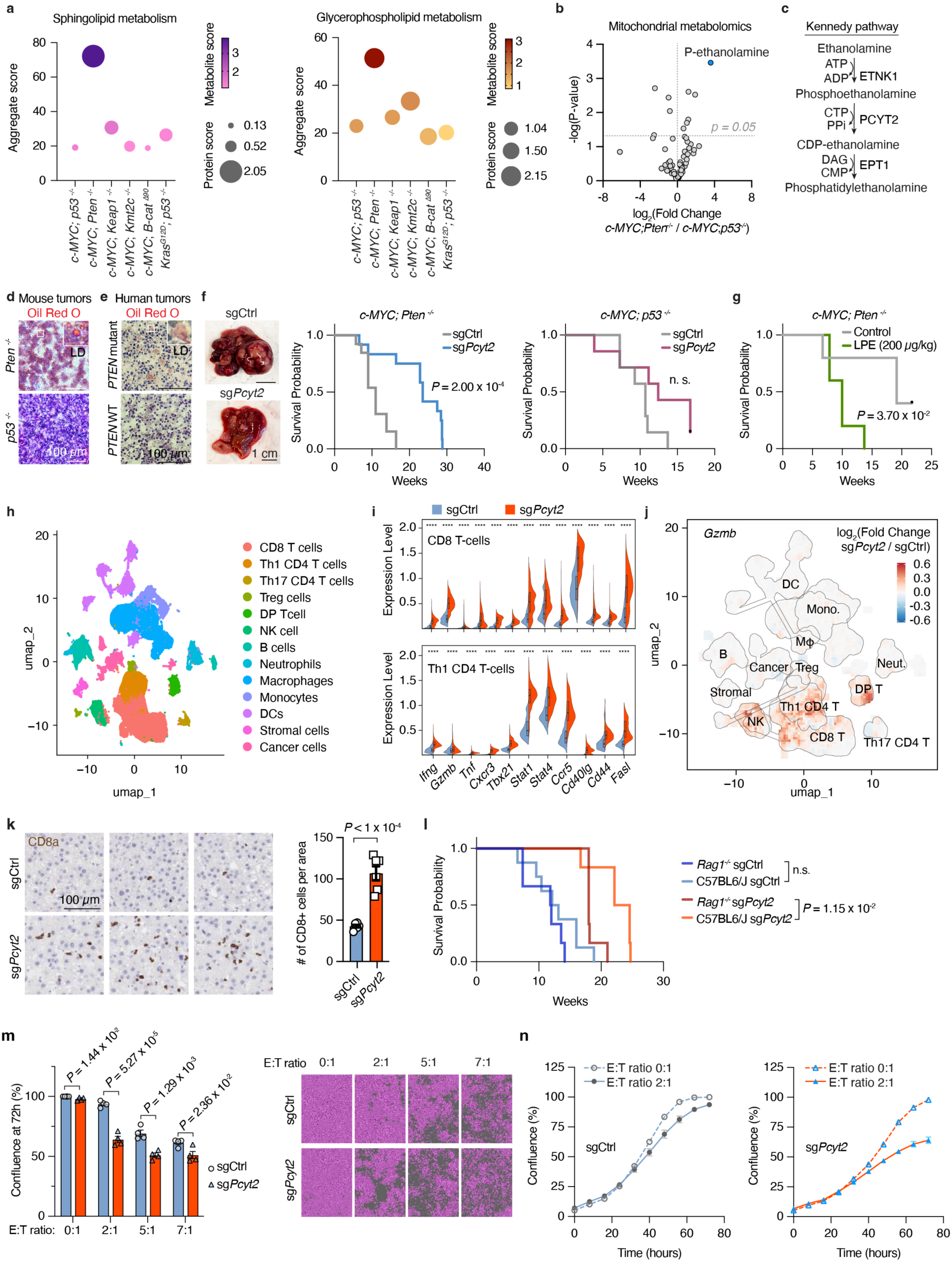
Essential role of PE synthesis in *PTEN*-null *c-MYC*-driven PLCs. a. Integrative network analysis showing enrichment of sphingolipid and glycerophospholipid metabolism in *c-MYC; Pten^-/-^* tumors as compared to other tumor models. b. Volcano plot for mitochondrial metabolomics comparing *c-MYC; Pten^-/-^* and *c-MYC; p53^-/-^* tumors (log_2_-scale) highlighting the enrichment of P-ethanolamine in *c-MYC; Pten^-/-^* tumors. -log *P*-value was calculated by Student’s t-test. Confidence interval is 95% c. Kennedy pathway for PE synthesis. ATP: adenosine triphosphate, ADP: adenosine diphosphate, CTP: cytidine triphosphate, PPi: inorganic pyrophosphate, DAG: diacylglycerol, CMP: cytidine monophosphate. d. Mouse *c-MYC; Pten^-/-^* and *c-MYC; p53^-/-^* tumors stained for Oil Red O and hematoxylin (counterstain, purple). Inset shows 4X magnified view of boxed area. LD :lipid droplet. Scale bar is 100 *μ*m. e. Human PLC tumors with and without *PTEN* mutations stained for Oil Red O and hematoxylin (counterstain). Inset shows 4X magnified view of boxed area. LD: lipid droplet. f. Images of *c-MYC; Pten^-/-^* tumors, both control (sgCtrl) and *Pcyt2*-depleted (sg*Pcyt2*). Survival curve for control and *Pcyt2*-depleted *c-MYC; Pten^-/-^* and *c-MYC; p53^-/-^* tumor bearing mice. Curve comparison by Mantel-Cox test, confidence interval 95%. g. Survival curve for vehicle control- and LPE (200 *μ*g/kg)-administered *c-MYC; Pten^-/-^* tumor bearing mice. Curve comparison by Mantel-Cox test, confidence interval 95%. h. UMAP plot showing dimensional reduction of cell types identified in tumors by sc-RNA analysis. Colors indicate cluster affiliation, i.e. cell types. i. Violin plots showing expression levels of immune cell activation markers in CD8 (top) and Th1 CD4 T-cells (bottom). Comparison bt Student’s t-test. ****, p < 0.0001. j. UMAP plot showing fold change (log_2_ scale) of *Gzmb* (right) expression across cell types, in *Pcyt2*-depleted (sg*Pcyt2*) vs. control (sgCtrl) tumors. k. IHC for CD8a in tumor sections from *Pcyt2*-depleted (sg*Pcyt2*) or control (sgCtrl) tumors. Bar plot shows quantification of number of CD8a^+^ cells per area of each tumor section. Comparison by student’s t-test, confidence interval is 95% l. Survival curve for control (sgCtrl) and *Pcyt2*-depleted (sg*Pcyt2*) *c-MYC; Pten^-/-^* tumor bearing *Rag1^-/-^* and immunocompetent C57BL6/J WT mice. Curve comparison by Mantel-Cox test, CI 95%. n.s., not significant. m. Proliferation of ovalbumin (OVA)-expressing control (sgCtrl) and *Pcyt2*-depleted (sg*Pcyt2*) *c-MYC; Pten^-/-^* cells untreated or co-cultured with OT-I CD8^+^ T-cells. Effector CD8^+^ T-cell (E) and target (T) ratios set up in cocultures are indicated. Left, bar graph depicting confluence of indicated conditions at 72 h. Right, representative images of the cultures at 72 h timepoint. Pink pseudocolor mask demarcates the regions occupied by cell bodies. Gray is the unoccupied region. Data are mean ± SEM; n = 4 biological replicates. n. Proliferation of OVA-expressing control (sgCtrl, gray) and *Pcyt2*-depleted (sg*Pcyt2*, blue) *c-MYC; Pten^-/-^* cells across 72-hour timescale. Data points represent confluence percentage every 8 hours. Dashed line indicates E:T, 0:1 and solid line indicates E:T, 2:1.

To test whether PE synthesis is selectively required for tumor progression in *Pten*-deficient PLCs, we targeted ethanolamine-phosphate cytidylyltransferase (PCYT2), the enzyme catalyzing the second step of Kennedy pathway (Figure 3c), in both *c-MYC; Pten^-/-^* and *c-MYC; p53^-/-^* tumors (Figure S7c). *Pcyt2*-deficient *c-MYC; Pten^-/-^* tumor-bearing mice exhibited a marked extension in median survival (∼23 weeks vs. ∼11 weeks for controls), whereas *Pcyt2* loss had no significant effect on overall survival of *c-MYC; p53^-/-^* tumor bearing mice (Figure 3f), indicating a genotype-specific dependency. Notably, histological analyses of *Pcyt2*-deficient *c-MYC; Pten^-/-^* tumors revealed enlarged vacuole-like structures that stained positive for lipid droplets (Figure S7d,e), accompanied by elevated triacylglycerol and diacylglycerol species (Figure S7f), consistent with enhanced flux of lipids into TAG stores upon Kennedy pathway blockade. To assess whether increasing PE abundance impacts tumor outcome, we further administered lysophosphatidylethanolamine (LPE), a membrane-permeable PE precursor, to tumor-bearing mice. LPE treatment resulted in reduced survival of *c-MYC; Pten^-/-^* tumor-bearing mice (Figure 3g), consistent with a potential limiting role for PE in supporting tumor progression. Together, these findings establish PCYT2-mediated PE synthesis as a genotype-specific metabolic dependency in *Pten*-deficient PLC.

### Tumor PEs enable immune evasion of *c-MYC; Pten^-/-^* PLCs

To mechanistically understand how *Pcyt2* loss results in delayed tumor growth, we asked whether it alters the immune landscape of the tumor microenvironment. To address this, we performed single-cell RNA sequencing (scRNA-seq) on a total of 80,000 cells from 8 tumors (4 control and 4 *Pcyt2^-/-^)* (Figures 3h and S8a). Genotype-based stratification revealed a marked expansion of CD8^+^ and Th1 CD4^+^ T cells in *Pcyt2*-deficient tumors (Figure S8b,c), accompanied by increased expression of cytotoxic and activation-associated genes including *Ifng*, *Gzmb*, *Ccl5*, and *Ctsw* (Figures 3i,j and S8d). In line with this observation, expression of immune-related receptors, including *Tnfrsf1b*, *Ifngr1* and *H2-K1* (MHC-I), was significantly upregulated in *Pcyt2*-deficient cancer cells (Figure S8e). Immunohistochemical staining for CD8a confirmed the increased infiltration of CD8^+^ T cells in *Pcyt2^-/-^*tumors (Figure 3k), consistent with scRNA and immune profiling data (Figure S8b,c). Increased immune infiltration within tumors can result in enhanced cell-cell communication between cancer and immune cells via ligand receptor interactions, ultimately influencing overall tumor cellular landscape and cancer progression. To investigate intercellular communication dynamics within *Pten^-/-^* tumors, we applied CellChat ^34^, a computational tool to infer intercellular signaling networks from scRNA-seq data. CellChat analysis identified heightened immune-related signaling between *Pcyt2^-/-^* cancer cells and infiltrating immune cells compared to control cells (Figure S8f). Notably, *Pcyt2*-deficienct tumors exhibited a greater number of inferred interactions between MHC class I (MHC-I) on cancer cells and *Cd8* on T cells (Figure S8g,h). To determine whether this immune engagement contributes to the anti-tumor phenotype, we generated *Pcyt2*^⁻^*^/^*^⁻^ tumors in both immunocompetent (C57BL/6J) and immunodeficient (*Rag1*^⁻/⁻^) mice, which lack functional T and B cells. While *Pcyt2* loss strongly prolonged survival in the immunocompetent context, anti-tumor effects were partially abrogated in *Rag1*-null mice, indicating that the tumor-suppressive effect of PE synthesis blockade is at least partially dependent on adaptive immunity (Figure 3l). Consistently, in direct co-culture assays, OT-I^+^ CD8^+^ T cells suppressed growth of OVA-expressing *Pcyt2*-KO cancer cells more effectively than controls, indicating enhanced immune sensitivity (Figure 3m,n). Collectively, these findings suggest that PE accumulation enables PTEN-deficient tumors to evade immune-mediated clearance, establishing PE synthesis as a metabolic mechanism of immune evasion.

### PE-mediated immune evasion through TNFα signaling

We next sought to define the mechanism by which PE synthesis promotes immune evasion of *PTEN* null PLCs. Cytotoxic lymphocytes eliminate cancer cells through the release of cytokines such as IFNγ and TNFα, which activate pro-apoptotic signaling in target cells^35^. Since membrane lipid composition influences cytokine receptor abundance and function, we considered that altered membrane homeostasis might be involved in the phenotypes we observed. To test this, we knocked out out *Pcyt2* in a mouse liver cancer cell line derived from *c-MYC; Pten^-/-^* tumors (Figure S9a,b) and performed proteomic analysis of cell surface proteins. Among the top upregulated surface proteins in *Pcyt2*-deficient cells compared to controls was the immune receptor TNFR2 (gene name, *Tnfrsf1b*) (Figure 4a and Figure S9b,c). Flow cytometry further confirmed increased surface expression of TNFR2 in *Pcyt2-KO* cells relative to both unedited cells and those deficient for sphingolipid synthesis (*Sptlc1*-KO) (Figure 4b and Figure S9d). TNFα receptors are internalized upon ligand stimulation^36^, yet surface levels of TNFR2 remained higher in in *Pcyt2*-KO cells upon TNFα stimulation (Figure 4c). To independently test this observation, we performed a FACS-based CRISPR screen using a lipid metabolism-focused sgRNA library in *c-MYC; Pten^-/-^* cell line. *Pcyt2* again ranked among the top hits associated with TNFR2 surface expression under both basal and TNFα-stimulated conditions, confirming the role of PEs in regulating receptor levels (Figure 4d and Figure S9e). Although TNFR2 lacks a death domain and cannot directly activate pro-apoptotic signaling, stimulation of TNFR2 was shown to inhibit the pro-survival arm of TNFα signaling, thereby activating apoptotic cascade ^37^. Consistently, *Pcyt2*-deficient cells showed enhanced caspase cleavage and impaired proliferation in response to TNFα treatment (Figure 4e,f). In parallel, to determine whether TNFα signaling contributes to CD8^+^ T-cell-mediated killing of *Pten^-/-^* liver cancer cells in an unbiased way, we performed a CRISPR screen using metabolic sgRNA library that included TNFα signaling genes. In this co-culture screen, *Tnfrsf1a* (encoding TNFR1) ranked amongst the top hits whose loss conferred resistance to T-cell-mediated cytotoxicity, along with *Tap1* and *Tap2,* key mediators of MHC-I antigen presentation (Figures 4g and S9f). Collectively, these results suggest that TNFα signaling is a critical mediator of CD8^+^ T-cell-mediated killing of *Pten^-/-^*liver cancer cells.

**Figure 4.**
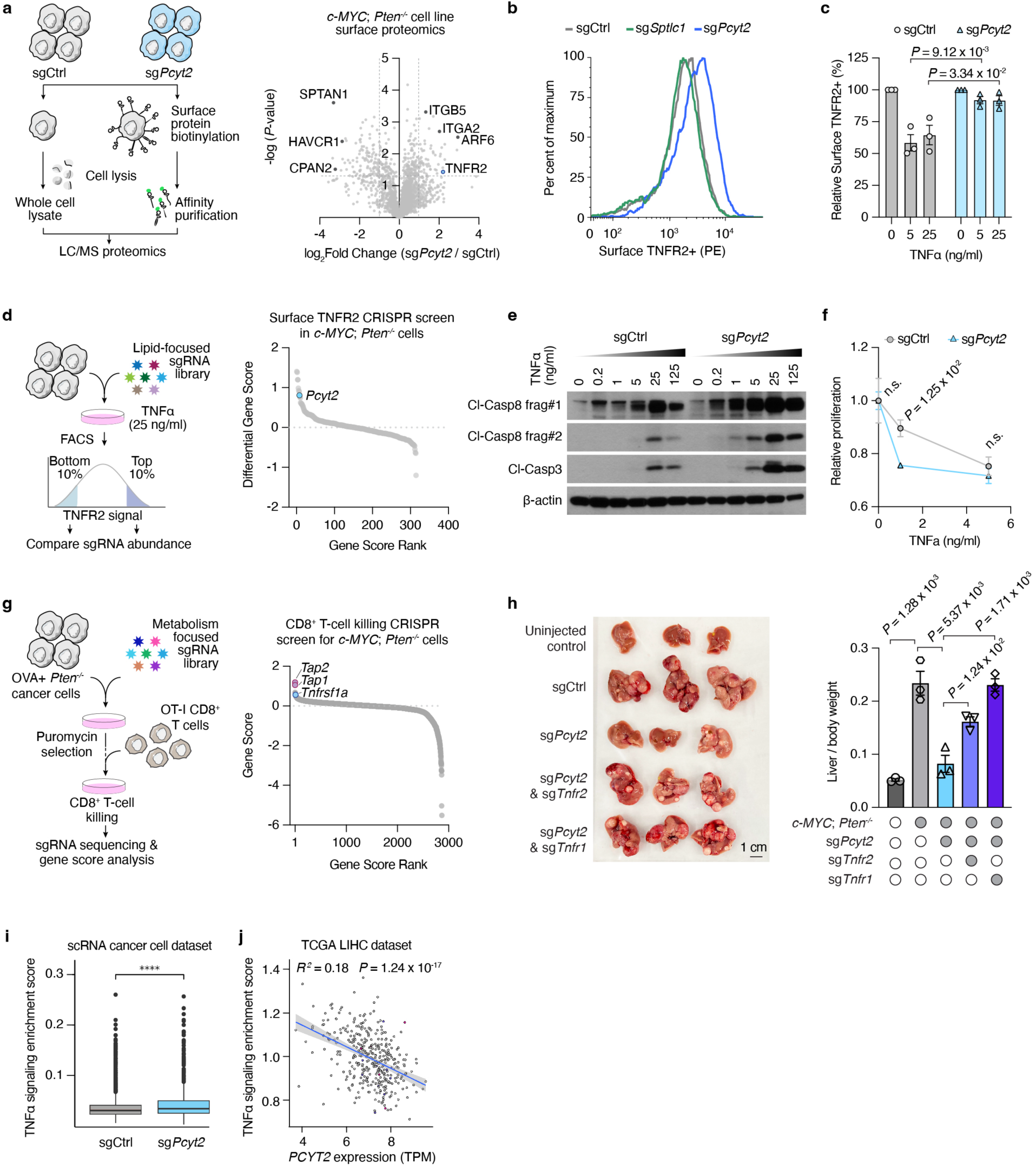
PE-mediated immune evasion through TNFα signaling. a. Control (sgCtrl) and *Pcyt2*-depleted (sg*Pcyt2*) *c-MYC; Pten^-/-^* cells incubated with a membrane-impermeable biotin tag. Cells were lysed and labeled proteins were affinity purified. Eluted protein lysates were analyzed by LC-MS. The log_2_-transformed fold change in membrane proteins between *Pcyt2*-depleted and control cells is displayed on volcano plot. b. Flow cytometry analysis of TNFR2 surface expression in sgControl, sg*Pcyt2* and sg*Spltc1* (used as a control for depletion of another membrane lipid class) *c-MYC; Pten^-/-^* cells treated with 25 ng/ml TNFα. PE: phycoerythrin. c. Flow cytometry analysis of TNFR2 surface expression in control and *Pcyt2*-depleted *c-MYC; Pten^-/-^* cells treated with 0, 5 and 25 ng/ml TNFα. Relative surface expression is plotted as per cent values normalized to untreated group. d. FACS-based screen to identify lipid metabolism genes that affect the stability of TNFR2 in the plasma membrane of *c-MYC; Pten^-/-^* cells. Live and transduced cells were incubated with phycoerythrin tagged antibody against TNFR2. High- and low-fluorescence populations were collected and amount of sgRNA was compared. Ranks of gene scores between TNFR2^hi^ and unsorted control population is plotted. *Pcyt2* was amongst the top 10 genes. e. Immunoblot for apoptosis induction in sgCtrl and sg*Pcyt2 c-MYC; Pten^-/-^* cells, untreated or treated with TNFα at varying concentrations for 16 hours. f. Proliferation of control (sgCtrl) and *Pcyt2*-depleted (sg*Pcyt2*) *c-MYC; Pten^-/-^* cells treated with the indicated concentration of TNFα for 16h. Data are mean ± SEM; n = 3 biological replicates. g. CD8^+^ T-cell mediated killing CRISPR screen for *c-MYC; Pten^-/-^* cells to identify cancer cell-intrinsic genes that affect cytotoxicity. *Tap1* and *Tap2* are components in MHC-I antigen presentation pathway, *Tnfrsf1a* encodes for TNFR1. These genes were amongst the top 10 hits. h. *Pcyt2*-single, *Pcyt2*; *Tnfr2*-double knockout and *Pcyt2*; *Tnfr1* liver tumors generated via HDTVi. Control tumors and uninjected healthy liver control are also included. Tumors were removed 9 weeks post-injection. Liver to body weight ratios are plotted as a reporter of tumor burden. Student’s t-test for comparison with 95% confidence interval. Data are mean ± SEM; n = 3 biological replicates. i. TNFα signaling enrichment score in control (sgCtrl) and *Pcyt2*-depleted (sg*Pcyt2*) *c-MYC; Pten^-/-^* cancer cell cluster from scRNA-seq dataset (from Figure 3). j. Correlation analysis of *PCYT2* expression and TNFα signaling enrichment score in TCGA liver hepatocellular carcinoma (LIHC) dataset. Each dot represents a patient. TPM: transcript per million.

To formally test whether TNFα receptor signaling mediates tumor growth defects observed upon *Pcyt2*-loss *in vivo*, we generated *Tnfr2^-/-^* and *Pcyt2^-/-^* as well as *Tnfr1^-/-^* and *Pcyt2^-/-^* double knockout tumors using HDTVi. While *Pcyt2*-KO tumors were significantly smaller than controls, co-deletion of *Tnfr2* or *Tnfr1* restored tumor growth (Figure 4h). Consistent with enhanced cytokine sensitivity of *Pcyt2*-deficient cancer cells, TNFα signaling signature was significantly elevated in *Pcyt2*-KO cancer cells within our scRNA-seq cancer cell dataset (Figure 4i). Finally, analysis of human liver hepatocellular carcinoma (LIHC) samples from TCGA showed a significant inverse correlation between PCYT2 expression and TNFα signaling enrichment score (Figure 4j). These findings identify PE synthesis as a regulator of TNFα receptor signaling and TNFα-driven immune surveillance in *PTEN*-null liver tumors.

## Discussion

Here, we utilized a cancer cell–specific organellar profiling platform for *in vivo* analysis of mitochondrial metabolites and proteins at cell-type and subcellular resolution, enabling the capture of metabolic states during transformation. By using mitochondria as a readout, this approach provides insight into cellular metabolic features in defined tumor cell populations and complements emerging spatial metabolomics tools. Extending this platform to other organelles such as lysosomes, peroxisomes, and the endoplasmic reticulum may reveal additional compartments through which tumor genotype shapes metabolism. A key insight from our study is that oncogene identity dictates mitochondrial metabolic programming and creates genotype-specific liabilities. In particular, *Kras*-driven tumors exhibited selective dependence on creatine metabolism during early tumor growth, likely reflecting the need for phosphocreatine-mediated energy buffering in the face of limited ATP production. This resembles muscle energetics, where glycolysis and OXPHOS are used to charge creatine pools, which in turn rapidly regenerate ATP during energy intensive periods ^38^. While creatine metabolism has been implicated in metastasis and hypoxia adaptation ^39, 40^, its role in early tumor growth has not been clear. Recent *in vivo* flux analyses confirm that solid tumors, including *Kras*-mutant models, produce ATP at lower rates than healthy tissue ^41^, supporting a model in which energy buffering becomes essential for sustaining proliferation of some cancer cells. In this setting, the creatine system may function as a critical energetic bridge, allowing tumors to buffer ATP fluctuations. Whether this requirement generalizes to other *Kras*-driven tumors including lung and pancreatic adenocarcinomas remains to be determined. In contrast, *c-MYC*-driven tumors depended on mitochondrial translation and OXPHOS, a recurrent metabolic phenotype across *c-MYC*-amplified cancers^42^ that may represent a tractable liability and potential biomarker for response to ETC inhibitors such as IACS-10759.

Beyond energy metabolism, our study highlights how cell-intrinsic lipid composition can regulate tumor– immune interactions. PE synthesis, selectively elevated in *PTEN*-null tumors, promoted immune evasion by suppressing TNFα sensitivity. Although TNF signaling typically supports NFκB-mediated survival, prior work has shown that it can also potentiate apoptosis via TRAF2 degradation ^37^. Consistent with this model, *Pcyt2*-deficient cells exhibited enhanced TNFα-induced caspase activation and increased susceptibility to T cell–mediated cytotoxicity. More broadly, changes in membrane phospholipid composition may affect lipid raft architecture and modulate receptor trafficking and cytokine responsiveness. Our findings add to a growing body of evidence that distinct lipid species including phosphatidylethanolamines, phosphatidylcholines, and sphingolipids ^43^ act as regulatory elements in signal transduction, and suggest that their therapeutic targeting may yield immunomodulatory benefits. The composition and functional impact of membrane lipids likely vary across oncogenic contexts, and our findings provide a proof-of-principle that certain tumor genotypes may be more vulnerable to interventions that target lipid metabolism. These therapies may include those modulating lipid availability through small molecule inhibitors or dietary interventions. Future studies systematically linking oncogenic alterations to membrane lipid metabolism across tumor types will be critical for uncovering generalizable principles and identifying context-dependent anti-cancer therapies.

## Acknowledgements

We thank all members of the Birsoy Lab for their feedback and suggestions. We also thank members of the Rockefeller University Proteomics Resource Center, the Flow Cytometry Resource Center for their technical support. G.U. is a Damon Runyon Fellow supported by the Damon Runyon Cancer Research Foundation (DRG-2431-21) and NIH/NCI 1K99CA286722-01. K.B. an Investigator of the Chan Zuckerberg Biohub New York; and supported by the NIH/NCI (1R01CA273233-01), NIH/NCI (5U54CA261701), a Mark Foundation Emerging Leader Award and Black Family Metastasis Research Center. N.D. is a William E. Ford Graduate Fellow at Rockefeller University. A.T. acknowledges the support of an Emmy Noether Award from the German Research Foundation (DFG, 467788900) and the Ministry of Culture and Science of the State of North Rhine-Westphalia (NRW-Nachwuchsgruppenprogramm). A.T. acknowledges the support of an ERC starting grant (METATARGET, 101078355). A.T. holds the Peter Hans Hofschneider endowed Professorship of Molecular Medicine from the Stiftung Experimentelle Biomedizin. The work of ISAS (A.T. and K.W.S) was supported by the “Ministerium für Kultur und Wissenschaft des Landes Nordrhein-Westfalen” and “Der Regierende Bürgermeister von Berlin, Senatskanzlei Wissenschaft und Forschung.” The work of K.W.S. was further supported by Federal Ministry of Education and Research (Bundesministerium für Bildung und Forschung, BMBF) under the funding reference 161L0271.

## Author contributions

K.B. and G.U. conceived the project. K.B. and G.U. wrote the manuscript with contributions from all authors. K.B. and G.U. designed experiments. G.U. ran most of the experiments with help from K.W., R.D., R.E., B.U. and N.D. A.M. performed most bioinformatics analyses including proteomics, metabolomics, integrated network analysis, bulk RNA-seq and human ICC cohort survival analyses with input from S.F.T. V.S. analyzed the scRNA-seq data and generated plots with inputs from S.B. K.C. and G.P. conducted lipidomics analyses. A.F. K.W.S., F.C., A.S. and A.T. performed MALDI-MSI spatial metabolomics analyses. Z.C.T. and O.B. provided human patient samples and helped with histological analyses. I.K. and S.F.T. provided RGX-202 diet and contributed to diet experiments. A.L. and R.D. provided plasmid constructs for HDTVi and provided feedback for genetic PLC models. K.C. and G.P. conducted lipidomic analysis. M.I.V, H.M. and E.K. performed polar metabolite profiling by LC/MS.

## METHODS

### Lead Contact

Further information and requests for resources and reagents should be directed to and will be fulfilled by the Lead Contact, Kıvanç Birsoy

### Materials Availability

All unique reagents generated in this study are available from the lead contact upon request.

### Data and code availability

Any additional information required to reanalyze the data reported in this paper is available from the lead contact upon request.

### Animal Studies

All animal studies were performed according to a protocol approved by the Institutional Animal Care and Use Committee (IACUC) at Rockefeller University. Animals were housed in ventilated caging on a standard light-dark cycle with food and water *ad libitum*. The wild type mouse strain C57BL/6J was obtained from the Jackson Laboratory (000664). To generate CMV-Cre;mito-tag mice, CMV-Cre male mice were purchased from Jackson Laboratory (006054) and crossed to mito-tag floxed female mice. Progeny was genotyped for the presence of mito-tag transgene and Cre recombinase.

### Human samples

All operatively resected tumors were collected after written patient consent and in accordance with the institutional review board approved protocols of Memorial Sloan Kettering Cancer Center (16-1683).

### Cell Lines, Compounds and Constructs

*c-MYC; Pten^-/-^* cell line was derived from PLC tumors generated by hydrodynamic tail vein injection. PLC tumor cells were dissociated using collagenase I (Thermo Scientific) and DNase I (Roche) enzyme mix in HBSS containing Ca^2+^ and Mg^2+^ (Gibco). Next, cells were washed in PBS and resuspended in DMEM/F12 (1:1) media (Gibco, 11330-032) supplemented with 10% fetal bovine serum and 1% penicillin and streptomycin; then plated in collagen coated flasks. Cells were maintained at 37°C, 21% O_2_ and 5% CO_2_This cell line was verified to be mycoplasma contamination-free.

### Generation of Knockout and Overexpression Lines

Knockout cells were generated with CRISPR/Cas9 method. Forward and reverse oligos targeting *Pcyt2*, or intergenic control were annealed and ligated into *Bsm*BI-linearized pLentiCRISPR v2 vector. For generation of knockout cells, VSV-G and Delta-VPR lentiviral packaging vectors were simultaneously transfected into HEK293T cells along with plentiCRISPR v2 vector expressing Cas9 and the gene-targeting sgRNA, using XtremeGene9 transfection reagent (Roche). Similarly, for overexpression of cytoplasmic OVA (cOVA), pLV-EF1a-IRES-Blast vector was transfected along with lentiviral packaging vectors VSV-G and Delta-VPR. 60 h post-transfection, the supernatant was collected after passing through a 0.45 *μ*m syringe filter. For transduction, 1 x 10^5^ cells were plated in 6-well plates containing 4 *μ*g/mL polybrene and virus, and then spin-infected by centrifugation at 2,200 rpm for 80 minutes. Mixed population knockouts were selected with puromycin; cOVA overexpressing cells were selected with blasticidin. Knockout efficiency was assessed via immunoblotting. OVA expression (presentation of SIINFEKL peptide by MHC-I) was assessed by flow cytometry using PE-conjugated H2-Kb bound SIINFEKL (Biolegend, #141603).

### Generation of cell-type specific mito-tag constructs

pCMV_SB13, pX330_p53sgRNA, pT3-EF1a-B-catΔ90 and pT3-EF1a-MYC-v5 construct was gifted from Lujambio laboratory at Mount Sinai. We modified the pT3-EF1a-MYC-v5 vector by cloning IRES-mito-Luciferase downstream of *c-MYC* coding sequence. Briefly, we linearized the pT3 vector with *Eco*RV digestion, then cloned two overlapping synthetic gene fragments for ‘IRES-3xHA-Luciferase-OMP25 localization signal’ (synthesized by Twist Bioscience) into the backbone by Gibson assembly. Of note, *Eco*RV digestion cleaves from within *c-MYC* coding sequence; thus, to re-assemble it back into pT3 vector, we amplified it from the original vector with 20-bp homologous arms and added into Gibson reaction along with mito-tag gene fragments. To generate Kras-IRES-mito-luciferase-tag plasmid, we replaced *c-MYC* coding sequence with *Kras^G12D^*, which we amplified from Caggs-Kras-IRES-EGFP construct (a gift from Lowe lab at MSKCC) with 20-bp homologous arms, then cloned via Gibson assembly. For sorting *Kras*-driven PLC cells via FACS, we followed a similar Gibson assembly strategy using ‘IRES-3xHA-tagRFP-OMP25 localization signal’ gene fragments that were then cloned into pT3-Kras backbone post-*Eco*RV digestion. Of note, this backbone still contains intact loxP sites as the original vector.

To generate cancer cell-specific gene depletion by CRISPR-Cas9 system for HDTVi, we constructed additional vectors. pT3-EF1a-c-MYC (without loxP sites, gifted by Lowe lab) was used the backbone. Following a similar cloning strategy, we inserted mito-luciferase-tag into this vector. Next, we cloned human U6 promoter and sgRNA scaffold from pLentiCRISPR_v2opti plasmid upstream of EF1a promoter. In addition, upstream of hU6 promoter, we inserted a second sgRNA site driven by mouse U6 promoter. mU6 driven site contained either the sgRNA of the tumor suppressor of interest (i.e. sg*Trp53* or sg*Pten* ) or a second sgRNA for the gene of interest. hU6 promoter driven site is amenable to standard sgRNA cloning by *Bsm*BI digestion as described previously for pLentiCRISPR backbones. Of note, spacer sequence dropped post-*Bsm*BI digestion was shortened to 580-bp at length. Lastly, to make gene depletion in *Kras* tumor background, we replaced *Kras^G12D^* coding sequence with *c-MYC* as described above.

Tumor suppressor sgRNAs were cloned into pX330 by linearizing the vector by *Bbs*I and ligating the annealed sgRNA oligos into the backbone. All sgRNA sequences are listed below:

*Trp53*: GCCTCGAGCTCCCTCTGAGCC, *Pten*: GAGATCGTTAGCAGAAACAAA, *Kmt2c*: GAATCAGTGCCAACCAATGGC, *Keap1*: GCTTATTGAGTTCGCCTACA, *Pcyt2*: GTGTCCACCACAGACCTCGT, , *Gatm*-sg1: GAAGCGTGAGCGCCATGCCA, Gatm-sg2: GCGAGATTATAGAAGCACCCA, *Mrps22*: GACACAGGCACAGTTGGAAG, sgControl (intergenic): GTGGGAACAGAGATAAGAAG, *Tnfr1*: GCAGCAGGCCAGGCACGGTG, *Tnfr2*: GTGCGGCCCTGGCTTCGGAG, *Sptlc1*: gAGGAAGAACTGATTGAAGAG

### Hydrodynamic tail vein injections

We prepared a plasmid mix containing 6.5 μg of pT3, 1.3 μg of pCMV-SB13 transposase, and 26 μg of pX330 expressing Cas9 cDNA and tumor suppressor sgRNA in sterile 0.9% NaCl solution. We then injected six to seven-week-old female mice through the lateral tail vein with a volume of plasmid/saline mix corresponding to 10% of body weight within 5–7 s as described previously ^15^. Tumor progression was monitored weekly by bioluminescence-based in vivo imaging system (IVIS). Total luminescence flux was measured, normalized to 3 days post-injection and plotted.

### Mitochondrial immunopurification from tissues

Mitochondrial immunopurification was performed as previously described with modifications ^20^. All procedures were performed on ice. The healthy liver or tumor tissue was excised with the punch biopsy tool (Integra, 4mm) and was immediately rinsed three times in ice-cold KPBS. Two pieces of liver tissue (∼50 mg total) were homogenized in 1 mL of KPBS (136 mM KCl, 10 mM KH_2_PO_4_, pH 7.25, in Optima LC/MS water) with a pestle attached to the mixer to rotate at 220 rpm in a 4 °C cold room for 30 seconds or until the tissue is fully homogenized. The homogenate was transferred to a low protein binding microfuge tube (Eppendorf), then spun down at 1,000 × g for 2 min at 4 °C. Ten microliters of the supernatant was taken as input sample and was extracted in the appropriate extraction buffer depending on the downstream analysis, namely metabolite or protein extraction. The supernatant was subjected to immunopurification with prewashed (in KPBS three times) anti-HA beads for 5 min on a rotator at 4°C, followed by three rounds of washing in KPBS to remove unbound material. In the final wash, 50% of the suspension of beads was separated for protein extraction and immunoblotting by aspirating the KPBS and then mixing the beads with 50 μL of Triton lysis buffer (50 mM Tris-HCl at pH 7.4, 150 mM NaCl, 1 mM EDTA, 1% Triton X-100, and Roche cOmplete EDTA-free protease inhibitor cocktail). The remainder 50% of the beads were then extracted in 80 % methanol containing ^15^N and ^13^C fully-labeled amino acid standards (MSK-A2-1.2, Cambridge Isotope Laboratories, Inc). 5 μL of input and immunopurified protein samples were taken and diluted in 45 μL Triton lysis buffer samples to be used for immunoblotting.

### Metabolite profiling by LC/MS

Polar metabolites were extracted in 80 % methanol containing ^15^N and ^13^C fully-labeled amino acid standards (MSK-A2-1.2, Cambridge Isotope Laboratories, Inc). Extracts were rotated end-to-end for 10 min at 4°C, spun at 19,000 g to remove insoluble cell debris, and stored at -80°C until liquid chromatography-mass spectrometry analysis (LC-MS). Then, LC-MS was performed as previously described ^44^. Relative metabolite abundances were quantified using XCalibur QualBrowser 2.2 and Skyline Targeted Mass Spec Environment ^45^ using a 5 ppm mass tolerance and a pooled-libraries of metabolite standards to verify metabolite identity. Relative metabolite levels were reported by normalizing to internal metabolite controls, NAD for mito-IP and tryptophan for whole tumor samples.

For whole tumor profiling, tissue samples were homogenized as described in mitochondrial immunopurification section, spun and metabolite extraction was performed immediately on homogenates.

### In vivo [^13^C_6_]-Arginine tracing

*KrasG12D; p53^-/-^* tumor-bearing mice were administered 20 *μ*L of 25 *μ*M [^13^C_6_]-Arginine solution in PBS (Cambridge Isotope Laboratories, CLM-2265-H-0.05) via the portal vein injection. Tumor and adjacent healthy liver tissue were harvested an hour post-injection and processed for the LC-MS analysis as indicated above. Per cent labeling of [^13^C_1_]-guanidinoacetate was reported.

### TMT proteomics analysis for mitochondrial proteome

Mitochondria were purified by mitochondrial immunopurification as described above. Proteins in elution buffer were reduced by dithiothreitol (DTT) and alkylated by iodoacetamide (IAA) followed by overnight ice-cold acetone precipitation Protein pellets were digested with both LysC and Trypsin in 200mM 4-(2-Hydroxyethyl)-1-piperazinepropanesulfonic acid (EPPS) buffer, pH 8.5. Resulting peptides were labeled using TMTPro 16-plex labeling reagents as described by vendor, and relative protein abundance across samples as well as labeling efficiency was evaluated. TMT-labelled peptides were mixed based on validation experiment and purified using reversed phase (RP) solid phase extraction. TMT peptides were fractionated through high pH RP fractionation as described by vendor (part# 84868, Thermo Fisher Scientific) and the resulting 8 fractions were analyzed by LC-MS/MS (EasyLC1200 coupled to a Fusion Lumos mass spectrometer, Thermo Fisher Scientific). The mass spectrometer was operated in DDA-MS2 High/High mode Spectra were searched against the mouse proteome concatenated with common contaminants, filtered using a 1% FDR and quantitated using Proteome Discoverer v.2.5 w/Sequest HT (Thermo Fisher Scientific). Data analysis was performed using Perseus v.1.6 ^46^.

Quantitation was performed using reporter ions from fragment spectra, requiring a spectral purity of 75%. Protein abundance values were log_2_-transformed. Proteins that were detected in 3 out of 3 replicates in at least one treatment group were kept and all other proteins were filtered out. Any missing abundance values were replaced by imputation.

### Lipidomics analysis by LC/MS

Liver tissue samples (∼20mg) were placed in 2 mL reinforced microtubes containing 2.8 mm ceramic beads (4-6 beads per sample) (Omni Cat#.19-628) and 2:2:1 methanol:acetonitrile:water (MAH; LC-MS Grade) at 40 *μ*L per mg of tissue was added to the tube. Samples were immediately homogenized using the Omni Bead Ruptor Elite Homogenizer (speed: 6 m/s; 2 cycles, each cycle run time: 20 seconds) equipped with a cooling hood attachment and the Omni BR-Cryo Cooling Unit at medium flow. Post-homogenization, samples were transferred into prechilled 1.7 mL microfuge tubes and stored at -20°C for 2 hours. After that, they were spun at maximum speed (> 12,800 rcf) for 10 minutes at 4°C. The supernatant was then transferred carefully to new microfuge tubes without disturbing the pellet (MAH portion, stored at -80°C). Pellets were resuspended in 40 *μ*L isopropanol (IPA; LC-MS Grade) per mg tissue, mixed thoroughly, and homogenized as previously described. Following identical incubation and centrifugation steps, the supernatant was transferred into separate microfuge tubes. Extracts for lipidomics were prepared by mixing MAH and IPA portions in a 1:1 ratio. The samples were stored at -80°C until LC/MS analysis.

Ultra-high performance liquid chromatography coupled with mass spectrometry (UHPLC/MS) analyses were performed using a Thermo Scientific Vanquish Flex UHPLC system, interfaced with a Thermo Scientific Orbitrap ID-X Mass Spectrometer. For the separation of lipids, an Acquity Premier HSS T3 column (2.1 x 100 mm, 1.8*μ*m) was utilized. The mobile-phase solvents consisted of solvent A = 10mM ammonium formate, 5 *μ*M medronic acid in 5:3:1 water:acetonitrile:2-propanol and solvent B = 10 mM ammonium formate in 1:9:90 water:acetonitrile:2-propanol. The column compartment temperature was maintained at 55°C and lipids were eluted using a linear gradient at a flow rate of 400 *μ*L/min as follows: 0 min, 15% B; 2.5 min, 50% B; 2.6 min, 57% B; 9 min, 70% B; 9.1 min, 93% B; 11 min, 96% B; 11.1, 100% B; 11.1-12min, 100% B; 12.2 min, 15% B; 12.2-16 min, 15% B. Data was acquired in positive and negative ion modes. The LC/MS data were then processed and analyzed using XCMS ^47^, CompoundDiscoverer, and Skyline ^45^.

### Immunoblotting

10-20mg tissue sample was washed in cold PBS, homogenized in microfuge tubes using polypropylene pestles and lysed in Triton lysis buffer (50 mM Tris-HCl at pH 7.4, 150 mM NaCl, 1 mM EDTA, 1% Triton X-100, and Roche cOmplete EDTA-free protease inhibitor cocktail). Lysates were sonicated, centrifuged at 1,000 g, and supernatant was collected as the protein lysate. Total protein quantified using BCA Protein Assay Kit (Thermo Fisher) with bovine serum albumin as a protein standard. Protein samples were resolved on 10%–20% SDS-PAGE gels (Novex, ThermoFisher) and analyzed by standard immunoblotting procedure. Briefly, PVDF membranes (Milipore) were incubated with primary antibodies at 4°C overnight. After washing off the primary antibodies in Tris-buffered saline/ 0.1% Tween-20 (TBS-T), secondary antibody incubation was performed at room temperature for 1h. HRP-linked IgG secondary antibodies including anti-mouse (Cell Signaling Technology, 7076) and anti-rabbit (Cell Signaling Technology, 7074) were used at 1:3,000 dilution. Blots were developed by ECL Chemiluminescent detection system (Perkin Elmer LLC) and film exposure. SRX-101A Film Processor (Konica Minolta) and Premium autoradiography Films (Thomas Scientific) were used for developing.

Primary antibodies used are Citrate Synthase (Cell Signaling Technology, 14309S), VDAC (Cell Signaling Technology, 4661S), CISD1 (Cell Signaling Technology, 83775S), OGDH (Proteintech, 15212-1-AP), GAPDH (GeneTex, GTX627408), SLC25A12 (abcam, 200201), Beta-actin (GeneTex, GTX109639), Cathepsin C (CTSC, Santa Cruz Biotechnology sc-74590), MRPS35 (Proteintech, 16457-1-AP), MRPS23 (Proteintech, 18345-1-AP), MRPS22 (Proteintech, 10984-1-AP), MRPL3 (Proteintech, 16584-1-AP), NDUFS1 (Proteintech, 12444-1-AP), NDUFB8 (Cell Signaling Technology, 73951S), Total OXPHOS cocktail (abcam, 110411), GATM (Proteintech, 66322-1-Ig), KRAS (Proteintech, 12063-1-AP), Phospho Acetyl CoA Carboxylase (Ser79) Antibody (Cell Signaling Technology, 3661S), PCYT2 (Proteintech, 14827-1-AP), Beta-tubulin (GeneTex, GTX101279), cleaved caspase-8 (Cell Signaling Technology, 8592T), cleaved caspase-3 (Cell Signaling Technology, 9661T), PTEN (Protein, 22034-1-AP), E-cadherin (Cell Signaling Technology, 14472S). All primary antibodies were diluted 1:1000 for immunoblotting assays.

### Sorting PLC cells with FACS and RNA-seq

To isolate early and late-stage cancer cells from *Kras*-driven tumors, mito-tag construct was modified to replace Luciferase with tagRFP coding sequence. KRAS-mito-RFP tumor bearing mice were sacrificed via cervical dislocation; tumors were dissected and minced with sterile scalpels. Tumor cells were dissociated with collagenase I (Thermo Scientific) and DNase I (Roche) enzyme mix in HBSS containing Ca^2+^ and Mg^2+^ (Gibco). Next, cells were washed in PBS and resuspended in FACS buffer (PBS containing 1% BSA and 1mM EDTA). Then, they were passed through 70 *μ*m strainer (Falcon). For FACS, dissociated cells were stained DAPI for viability. Next, RFP+ DAPI-population was sorted directly into TRIzol-LS with Sony MA900. Then, total RNA was isolated following TRIzol-LS (Thermo Fisher Scientific) manufacturer’s manual. RNA concentrations were determined using Qubit 2.0 Fluorometer (Life Technologies); RNA integrity was checked using Agilent TapeStation 4200 (Agilent Technologies). RNA sequencing library was prepared using NEBNext Ultra RNA library kit for Illumina (NEB) following manufacturer’s manual. The sequencing libraries were clustered on a single lane of a flowcell. Afterwards, the flowcell was loaded on the Illumina HiSeq instrument (4000 or equivalent) according to manufacturer’s guidelines. The samples were sequenced using a 2x150bp Paired End (PE) configuration. For RNA-seq analysis, sequence and transcript coordinates for mouse genome (mm10 UCSC) and gene models were retrieved from the Bioconductor Bsgenome.Hsapiens.UCSC.mm10 (version 1.4.0) and TxDb.Hsapiens.UCSC.mm10.knownGene (version 3.4.0) Bioconductor libraries respectively. Transcript expressions were calculated using the Salmon quantification software ^48^ (version 0.8.2) and gene expression levels as transcripts per million (TPM). Normalization and rlog transformation of raw read counts in genes were performed using DESeq2 ^49^ (version 1.20.0). Genes significantly differentially expressed between conditions were identified using DESeq2 with a Benjamini Hochberg adjusted p-value cutoff of 0.05. Gene set enrichment was obtained for all genes significantly differentially expressed between conditions (absolute logFC > 0, adjusted p-value < 0.05) using the fisher test in the topGO Bioconductor package and ranked using the elim algorithm and functional annotation from the org.Mm.eg.db Bioconductor package (version 3.10).

### Immunofluorescence (IF) and Immunohistochemistry (IHC) analysis

Liver tissue was fixed in 10% formalin over 48 hours at room temperature, then dehydrated in 70% ethanol. Tissue was paraffinized and sectioned at 4-10 *μ*m thickness. After deparaffinization and rehydration, the tissue sections were immersed in Citrate-Based antigen unmasking solution (VectorLabs), and subjected to pressure cooker treatment for 15 min for antigen retrieval. The samples were subsequently immersed in blocking solution (3 mg/ml BSA in PBS) for 15 mins and then incubated with primary antibody diluted in blocking buffer 1 hour at RT. For the multiplexed immunofluorescence, the tissue slides were stained with corresponding Alexa Fluor-488, Alexa Fluor-555 conjugated secondary antibodies. The antibody-labelled slides were stained with 4′,6-diamidino-2-phenylindole (DAPI) as nuclear counterstain; then mounted using ProLong Gold antifade mountant (Molecular Probes). Imaging was performed using a Nikon A1R MP multiphoton microscope in confocal mode with a Nikon Plan Apo γ 60X/1.40 oil immersion objective. Line scan for colocalization analysis was conducted using ImageJ (Fiji version 2.14.0). Briefly, a straight line was drawn in multi-color image and fluorescence intensity was measured throughout the distance of the line. Data were imported to GraphPad Prism (v10) and plotted as x-y graphs, Distance (*μ*m vs. fluorescence intensity).

The following antibodies were used at 1:100 dilution for IF: HA-tag (Cell Signaling Technology, 2367S), Citrate Synthase (Cell Signaling Technology, 14309S), ATP5A1 (ThermoFisher Scientific, 439800), LAMP-1 (Developmental Studies Hybridoma Bank, 1D4B), Calreticulin (Cell Signaling Technology, 12238P), KRAS (Proteintech, 12063-1-AP), Ki67 (Proteintech, 27309-1-AP), CD45 (Proteintech, 201031-1-AP), CD31 (GeneTex, 20218), c-Myc (Abcam, ab32072).

For the IHC (Immunohistochemistry) detection, the slides were processed using an EnVision+Dual Link System-HRP kit (DAKO) according to the manufacturer’s protocol, and counterstained using haematoxylin; then mounted using VectaMount mounting solution (VectorLabs). Images were acquired using Echo Revolve microscope with Nikon 10X objective.

The following antibodies are used at 1:100 dilution in this study: CD8 alpha (Cell Signaling Technology, 98941), Phospho Acetyl CoA Carboxylase (Ser79) Antibody (Cell Signaling Technology, 3661S).

### IACS-10759 treatment

Three days post-tumors establishment via HDTVi, mice were treated with 5 mg/kg IACS-10759 (MedChem Express) through oral gavage. IACS-10759 was prepared by dissolving it in DMSO (10%), then in corn oil (90%). As control, 10% DMSO + 90% corn oil mix was given to a set of animals with tumors. All mice treated on a 2 days on / 2 days off regimen. Body weights were measured, and no significant loss was detected under this regimen. Tumor sized were monitored and recorded by IVIS measurements at 3 days (initial) and weekly measurements until 3 weeks final timepoint.

### Lysophosphatidylethanolamine (LPE) administration

Mice were injected with 200 *μ*g/kg every other day starting from the day of HDTV injection. Injection regimen was adapted from Xu et al. study ^50^. Control mice were injected with buffer per the same regimen.

### RGX-202 treatment

Tumor bearing mice mice were fed creatine metabolism inhibitor, RGX-202 at ∼650 mg/kg (calculated based on daily consumption) or standard chow diet (Purina 5001) *ad libitum*. Tumor progression was monitored by weekly IVIS measurements.

### Quantification and statistical analysis

Statistical analysis was performed with built-in statistics tools in Prism10 (GraphPad Software). Statistical tests, error bars and confidence intervals can be found in the figure legends. p-values for comparisons between groups are indicated in figures.

### Oil Red O staining

Liver tissue was dissected and rinsed in PBS, then fixed overnight in 4% parafolmaldeyde (PFA) at room temperature. Next day, PFA was removed, and tissue was stored in 30% sucrose (in PBS) overnight at 4 °C. For cryosectioning, tissue was embedded in O.C.T. (Fisher Scientific), frozen at -20°C, then 14 *μ*m thick sections were collected with a cryostat (Leica). After that, slides were air dried for 30 minutes, then fixed in 10% neutral buffered formalin (Sigma-Aldrich). Following fixation, slides were dipped in 60% isopropanol quickly, and then dipped in Oil Red O solution (0.18% in 60% isopropanol) for 15 minutes. Following staining, slides were dipped in 60% isopropanol and deionized water sequentially. As counterstain, slides were incubated in Mayer’s hematoxylin (VectorLabs) for three minutes, then rinsed in deionized water and mounted using VectaMount mounting solution (VectorLabs). Images were acquired using Echo Revolve microscope equipped with Nikon 10X objective.

For human patient tissues, cryosections were collected from flash frozen, unfixed specimen. The rest of the staining protocol was followed as described above.

### FACS-based CRISPR screen

*c-MYC; Pten^-/-^*cells (2.5 million) were infected with lipid focused sgRNA library ^43^ at an MOI of ∼0.7 and selected with puromycin. Following selection, an initial pool of 2.5 million cells was collected for genomic DNA extraction and transduced cells were cultured for another 24 hours. 50 million cells were collected and washed once with PBS. Cells were resuspended in 400 *μ*l FACS buffer (PBS, 1% BSA, 5 mM EDTA) containing 1:100 diluted PE-conjugated anti-CD120b (TNFR2) (Biolegend, #113405) and DAPI (viability marker), incubated at 4°C for 15 min. Then, cells were washed with 10 ml FACS buffer and spun at 500 g for 5 min at 4°C. Pellets were resuspended in FACS buffer. Top and bottom 10% PE+ live cells were sorted using Sony MA900. Genomic DNA extraction, PCR amplification and quantification of sgRNAs and gene score analysis was conducted.

### CD8+ T-cell isolation, expansion

The spleen from C57BL/6-Tg (TcraTcrb)1100Mjb/J (OT-1) mice were collected and dissociated on a 35-mm plate over ice using a scalpel. The tissue was resuspended in cold MACS buffer (PBS, 0.5% BSA and 2 mM EDTA) and passed through a 70-μm cell strainer. The plunger of a 3-ml syringe was used to push remaining tissue through the strainer, which was then rinsed with MACS buffer. Cells were centrifuged at room temperature for 5 min at 1,200 rpm. RBC lysis was performed by incubating cells in ACK lysis buffer (Thermo Scientific A1049201) at room temperature for 5 min. An equal volume of MACS buffer was added to the cell suspension and transferred through a 70-μm cell strainer into a new tube. Cells were centrifuged again at room temperature for 5 min at 1,200 rpm, resuspended in 1 ml of MACS buffer, and counted. CD8+ T cells were isolated from the cell suspension using negative selection isolation kits following the manufacturer’s protocols (Miltenyi Biotec 130-104-075). CD8^+^ T cells were cultured for overnight at a concentration of 10^6^ cells per ml in T cell medium (RPMI 1640 medium, 2 mM glutamine, 1 mM pyruvate, 10% FBS, 1% penicillin/streptomycin, 1× MEM amino acids (Thermo Scientific 11130051), 50 μM mercaptoethanol, pH ∼7.4) with 20 ng ml^−1^ IL-2. T cell activation was performed by culturing cells with 1 *μ*g/ml ovalbumin for 6 hours before co-culturing assays.

### CD8+ T-cell – cancer cell co-culture assays

To allow adherence, *c-MYC; Pten^-/-^* cells were plated in triplicate at a concentration of 1,000 cells per 100 *μ*l per well in a 96-well plate for 24h before the addition of CD8^+^ T-cells. One hundred microliters of activated CD8^+^ T-cells was immediately added to cancer cells with 5 ng ml^-1^ IL-2 supplementation, at the respective E:T ratios. *c-MYC; Pten^-/-^* cell death and proliferation was monitored over 3 days using incucyte.

### CD8^+^ T-cell killing CRISPR screen

*c-MYC; Pten^-/-^*cells (25 million) overexpressing cOVA were infected with mouse metabolism-focused sgRNA library^51^ at an MOI of ∼0.7 and selected with puromycin. Following selection, an initial pool of 25 million cells was collected for genomic DNA extraction and transduced cells were cultured for another 24 hours. Then, 5 million cells (0.5 million per 10 cm^2^ dish, a total of 10 dishes) were co-cultured with CD8^+^ T-cells isolated from OT-I mice at 2:1 E:T ratio (i.e. 1 million T cells per 0.5 million cancer cells per dish) in T-cell medium (RPMI 1640 medium, 2 mM glutamine, 1 mM pyruvate, 10% FBS, 1% penicillin/streptomycin, 1× MEM amino acids (Thermo Scientific 11130051), 50 μM mercaptoethanol, pH ∼7.4) with 5 ng ml−1 IL-2. For control, the same number of cancer cells were cultured in the medium. 72 hours later, untreated cells and those co-cultured with CD8^+^ T-cells were collected for genomic DNA extraction. Then, PCR amplification, quantification of sgRNAs and gene score analysis was conducted.

### Flow cytometry analysis

*c-MYC; Pten^-/-^* cells (500,000) were collected and washed once with PBS. Cells were resuspended in 50 μl FACS buffer containing 1:100 diluted PE-conjugated anti-CD120b (TNFR2) (Biolegend, #113405) and DAPI (viability marker), and incubated in the dark at 4 °C for 15 min. To wash cells, 1 ml of FACS buffer (PBS, 1% BSA, 5 mM EDTA) was added to each sample before centrifugation at 500g for 5 min at 4 °C. Pellets were resuspended in FACS buffer and immediately analysed on an Attune NxT (Thermo Scientific).

### Flow cytometry analysis of tumor-infiltrating immune cells

Tumors were excised and digested for 1 h at 37 °C with with 400 U ml^−1^ of collagenase D (Roche) and 0.2 μg ml^−1^ DNase I (Sigma). Digested tumors were filtered through 70-μm filters. Enrichment of hematopoietic cells was achieved by density gradient centrifugation with 40% / 80% Percoll (GE Healthcare Life Sciences) for 25 min at 2,500 rpm at 22 °C without breaks. The interphase containing hematopoietic cells was isolated and washed with phosphate-buffered EDTA (PBE). Red blood cell lysis was performed with ACK lysis buffer (GIBCO). Cells were incubated for 5 min with 1 μg ml^−1^ of anti-CD16/32 (2.4G2, BioXcell) on ice for 5 min. Cells were washed with PBS and stained with appropriate surface marker antibodies for CD45 (Biolegend, 103115), TCR-beta (Biolegend, 109228) and CD8a (Biolegend, 100707) for 20 min at 4 °C. Cells were then washed and resuspended in PBE. Samples were acquired on Attune NxT (Thermo Scientific).

### Membrane protein biotinylation and isolation

*c-MYC; Pten^-/-^*cells were plated 18 h before collection so that they would reach 80% confluency on the day of collection. After respective incubation times, all samples were collected at the same time. All steps were performed on ice or at 4 °C. Cells were washed 3 times with 10 ml ice-cold PBS. 10 ml of 20 mM EZ-Link Sulfo-NHS-LC-Biotin (Thermo Scientific 21335) was added to the cells. Plates were incubated at 4 °C for 30 min. Plates were washed once with 10 ml ice-cold TBS (pH 7.4) and then twice with ice-cold PBS. Cells were then scraped off the plate, collected, and centrifuged at 500g for 5 min at 4 °C. Cells were resuspended in RIPA buffer containing protease inhibitors and incubated on ice for 30 min. Lysates were sonicated for 5 min and then centrifuged at maximum speed for 10 min at 4 °C. Lysates were transferred to streptavidin beads (Thermo Scientific 88816) and incubated on a rotator at 4 °C for 1 h. Beads were briefly spun down, magnetized, and washed 3 times with 1 ml of PBS. Protein was eluted off of the beads by resuspending them in 50–100 μl of transmembrane buffer (10 mM Tris-HCl pH 7.4, 1 mM EDTA, 1% Triton X-100, 2% SDS, 0.1% CHAPS) with 1× SDS loading buffer (20% SDS) and boiling samples for 5 min at 95 °C. Beads were then magnetized and the lysate collected.

### Batch correction and clustering of metabolomics and proteomics data

We loaded raw data from mass spectrometry of metabolites from mitochondrial IP into R and merged it across all batches. To impute missing values, we used the ‘mice’ package ^52^ under the classification and regression tree (‘cart’) method over 5 cycles. Batch effects were then regressed out using the ‘ComBat’ package ^53^, correcting for batch while preserving genotype and restricting to parametric adjustments. This imputed and batch-corrected data was used for downstream analysis. The same approach was utilized to generate imputed and batch-corrected data from mitochondrial IP proteomics. We performed clustering both by PCA and by hierarchical clustering via the algorithm of Gruvaeus and Wainer as implemented by the ‘seriation’ package ^54, 55^.

### Non-negative matrix factorization of metabolomics data

We first Z-scored and base-e exponentiated the imputed and batch-corrected metabolomics data to produce a non-negative matrix. To estimate the optimal rank value for NMF decomposition, we swept across ranks 2-12 and ran 50 rounds of NMF per rank, selecting an initialization seed via independent component analysis. Based on this parameter sweep, a rank of 6 was identified as optimal. NMF was finally implemented ^56^ with this value and calculated over 200 runs with initialization seed defined by independent component analysis.

### Metabolic-proteomic network analysis

#### Computation of enrichment scores

In order to build networks linking metabolites to proteins, we first leveraged the well-annotated ConsensusPathDB ^57^resource to produce a dictionary of pathways and their matching metabolites and proteins. We filtered only to overlapping pathways that were annotated in both metabolites and proteins and converted protein names from human to mouse using the MGI homolog reference. For each tumor type, we computed the p-value and logFC of all proteins and metabolites compared to healthy controls based on the imputed and batch-corrected data and using the fast Wilcoxon rank sum test and auROC implementation in the ‘presto’ package ^58^. The logFC values for each metabolite and each protein were then ranked within each tumor type, resolving ties by the “dense” method to ensure no gaps in the order. These statistics were used to define an **enrichment score** for each pathway in each tumor type as follows: 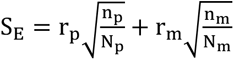, where r_p_, r_m_ are the logFC rank for proteins and metabolites in that pathway, respectively, n_p_, n_m_ are the number of proteins and metabolites in that pathway detected in the data, respectively, and N_p_, N_m_ are the total number of proteins and metabolites annotated in that pathway, respectively. In brief, this formula sums the ranked logFC of proteins and metabolites in a tumor type for given pathway, weighted by the detection rate of those proteins and metabolites in the immunopurification data.

#### Identification of enriched pathways

Given these S_E_ enrichment scores, we subsequently aimed to identify pathways that were specifically enriched or depleted in distinct tumor types. To accomplish this goal, we first calculated the score deviation, 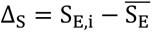, or the difference between the S_E_ enrichment score for one tumor and the mean S_E_ enrichment score for all tumor types. The score deviations were normalized to fractional deviations, 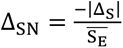, to account for the range of S values across pathways. We ranked the original scores as well as these normalized score deviations to generate two rankings, one of the scores themselves within a tumor type (R_S_) and one of their relative enrichment in that tumor type (R_Δ_). We aggregated these statistics together into a final **aggregate score**, 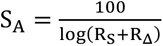, which accounts for both the extent to which a pathway is enriched relative to other pathways within a tumor and the extent to which a pathway is enriched in one tumor relative to other tumors. This aggregate score was used for downstream visualization.

#### Identification of depleted pathways

To compute depleted pathways, we instead filtered only to pathways where both the protein and metabolite mean logFC values were negative. We calculated score deviation Δ_S_ as defined above and computed a depletion score, 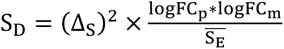, where logFC_p_, logFC_m_ are the mean logFC values for proteins and metabolites in the pathway for the tumor type and 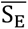 is the mean enrichment score across all tumor types. These depletion scores were normalized by dividing by the maximum S_D_ value within the tumor type and then used for downstream visualization.

#### Visualization of pairwise tumor-metabolite interactions

To finally plot pairwise interactions in the network that were enriched or depleted in a genotype-dependent manner, we first computed the pairwise Pearson correlation between each protein and each metabolite separately in each tumor type. We then filtered to the top five pathways by aggregate score S_A_ in each tumor type and computed the logFC value between the tumor type and healthy controls for all metabolites and proteins in those five pathways. To make comparisons between metabolites and proteins possible, we dense ranked these logFC values and then normalized them by dividing by the maximum rank, r_p_/ max:r_p_; and r_m_/ max(r_m_). We plotted these pairwise protein-metabolite interactions as a dotplot, where the x-axis is a protein, the y-axis is a metabolite, the size of the point is the product of the ranks r_p_ × r_m_, and the color is the Pearson correlation coefficient of the metabolite-protein pair.

### RNAseq analysis

We generated counts matrices from the raw fastqs by pseudoalignment against mm10 using salmon and loaded the resulting tables into R with tximport ^48^. Samples were normalized and variance-stabilized and differentially expressed genes were called through the limma-voom pipeline ^59^. The logFC and p-values computed by limma-voom for all genes were used as inputs for pathway analysis using the fGSEA implementation ^60^. We focused on pathways related to bioenergetics, manually identifying nine pathways annotated in Reactome, WikiPathways, and GO with relevance to energy homeostasis and metabolism and plotting the fGSEA random walks from these pathways directly.

### TNFa signature analysis

Bulk RNAseq data from TCGA: We normalized and variance-stabilized the raw counts data from TCGA via the limma-voom pipeline as described above. To evaluate TNFa signaling in each sample, we computed the GSEA enrichment score per sample for the HALLMARK_TNFA_SIGNALING_VIA_NFKB genelist using fGSEA.

Single cell data: We used AUCell to compute a per-cell integrated expression score over all genes in the HALLMARK_TNFA_SIGNALING_VIA_NFKB genelist, using the raw RNA counts as input data. To visualize these results, we plotted TNFa scores for each cell as boxplots grouped by “supercategories” of cluster identities. We also constructed a generalized linear model with a log-link Gamma distribution modeling the impact of Pcyt2 status, supercategory, and their interaction on TNFa signature score.

### Survival curve

To determine whether the expression of creatine metabolism genes correlated with survival in the intrahepatic cholangiocarcinoma tumor dataset, we focused on GATM, GAMT, CKB, CKM, CKMT1, and SLC6A8 as a clearly defined set of genes specifically involved in creatine metabolism. We used the ‘survival’ package ^61^ to fit individual Cox proportional-hazards models for each of these genes both at the transcriptional and at the proteomic level and used the coefficients from these models to produce a final weighted score for each patient representing the total incremental risk represented by each RNA and protein feature defined as the product of the expression and its coefficient. We ran a final Cox proportional-hazards model on the weighted scores and split patients into “High” or “Low” creatine metabolism groups based on whether their weighted score was greater than or less than the median value across the dataset.

### Single cell RNA sequencing data analysis

Single cell RNA sequencing was performed using Parse Evercode WT kit following manufacturer’s instructions. Four tumor bearing mice per genotype were included. Single cell datasets were assessed for data quality following the guidelines described previously ^62, 63^. Cells with more than 10% mitochondrial transcripts as well as cells that had fewer than 500 feature counts or expressed fewer than 100 genes were removed. After quality control (QC), Seurat (v5.0.1) ^64^ was used for normalization, graph-based clustering and differential expression analysis. Each dataset was normalized using SCTransform and the 2000 most variable genes were identified with SelectIntegrationFeatures. MAGIC imputation was conducted on normalized data to impute missing values and account for technical noise ^65^. RunPCA was implemented on the normalized datasets to identify the top 25 principal components (PCs) that were used for UMAP analysis and clustering. UMAP was calculated using the runUMAP function. Clustering was conducted by first constructing a nearest neighbor graph using the FindNeighbors function and then implementing the FindClusters function to perform clustering using the Louvain algorithm at a resolution of 1. Clusters were labeled in accordance with CD45 tumor infiltrating lymphocyte subtype signatures identified by Zheng et al. study ^66^. Differential expression analysis was conducted between groups using the FindMarkers function with the MAST method to evaluate differences within the transcriptome ^67^. Wilcoxon rank-sum tests to determine if gene expression was significant was conducted using the wilcox.test function in stats (v4.1.0) ^34^. Differential interactome modeling was conducted using the CellChat package (v2.1.0). Gene set enrichment analysis was conducted using EnrichR (v3.2) ^68^. Scores were calculated using method detailed previously. Pearson correlation coefficient test was calculated to evaluate the significance of the association. Analysis code is available on GitHub (https://github.com/Vyoming/Azenta-Project-30-1097416454.git).

### MALDI-MSI imaging and data analysis

#### Tissue preparation for liver staining and MALDI imaging analysis

Liver tissue was sectioned at 12*μ*m thickness using a Leica CM1860 cryotome. For MALDI imaging, tissues were thaw mounted onto ITO glass slides (70-100 ohms, Delta Technologies, USA), with serial sections collected on regular histology glass slides for staining protocols. Sectioning was performed at -16°C, with samples being stored at -80°C until analysis.

#### MALDI matrix and internal standard application

NEDC (N-naphthylethylenediamine dihydrochloride; 7mg/ml in 70:25:5 MeOH:ACN:H_2_O) was applied via a SunCollect Sunchrom sprayer (9-port dispenser version). Briefly, 20 layers were applied using a flow rate sequence of 1-8 layers at 15*μ*L/min and 8-20 layers at 20*μ*L/min at a speed of 620 mm/s. Gas pressure was set at 2.5 bar and a z-position of 35mm.

#### MALDI-MSI instrumentation

MSI was performed on an elevated pressure MALDI source (Spectroglyph LLC, Kenwick, WA, USA) coupled to a Thermo Q-Exactive HF mass spectrometer (Thermo Fisher Scientific, Bremen, Germany). A 349-nm Nd:YLF laser (Explorer One, Spectra Physics, CA) was operated with a repetition rate of 500Hz and a pulse energy of 1-2*μ*J. Imaging analysis was carried out using a spacing raster of 20x20*μ*m, with a laser spot size of ∼15*μ*m. For negative ion mode analysis, the ion funnel was operated with 85V_pp_ (HPF) and 75V_pp_ (LPF) using a 15% RF drive for both funnels for 720kHz and 825kHz, respectively. For positive ion mode analysis, the ion funnel was operated with 225V_pp_ (HPF) and 125V_pp_ (LPF) using a 35% (HPF) and 30% (LPF) drive for 720kHz and 825kHz, respectively. Source operating pressure was maintained at 6.5 Torr.

The mass spectrometer was operated in positive and negative mode with AGC off and at 250ms injection time. Mass range was set at *m/z* 50-750 with resolution set at 120,000 at *m/z* 200. Instrument calibration was carried out using the ESI interface with Pierce Negative Calmix solution spiked with metabolite standards.

#### Tissue preparation for LC-MS analysis

Liver tissue was sectioned and collected for bulk LC-MS analysis by sectioning in the cryotome. An organic solvent-based protein precipitation was used to do metabolite extraction, previously described (Kasarla et al 2024). Briefly, 350*μ*L of extraction solvent (40:40:20 v/v, MeOH:ACN:H_2_O) was added to the samples, and homogenized using a Pellet Pestle (Kontes, USA) followed by sonication using Bioruptor Pico (Diagenode, USA) at a temperature of 4°C and seven cycles (30s on/ 30s off) with glass beads to facilitate the homogenization. Samples were then vortexed at 4°C for 10 minutes and left for incubation at -80°C overnight for protein precipitation. Centrifugation then followed for 15,000 RPM for 30 minutes at 4°C. The supernatant was separated from the pellet and dried under nitrogen gas at room temperature. When dried, samples were reconstituted in ACN:H_2_O (70:30 v/v) vortexed for 10 minutes at 4°C. After a further 2-hour incubation at -80°C, the sample extracts were centrifuged at 13000 rpm and 4°C for 30 minutes and the supernatant was transferred to LC-MS vials.

#### LC-MS/MS methodology for bulk tissue analysis

Measurements were performed using a Vanquish Duo UHPLC-system (Thermo Fisher Scientific, Waltham, MA, USA) equipped with a dual pump, an autosampler, and a thermostatic column compartment with the Atlantis Premier BEH Z-HILIC column. The mobile phases used were 100 % water with 10 mM ammonium acetate and 0.1 % ammonium hydroxide, pH 8.5 as a mobile phase A and 100 % ACN as a mobile phase B with a flow rate of 0.4 mL/min. The elution gradient conditions were as follows: 95 % B, 0–2 min; 95–80 % B, 2–7.7 min; 80–70 % B, 7.7–9.5 min; 70–10 % B, 9.5–10 min followed by a 2-minute hold; 10–30 % B, 12–16 min; 30–95% B 16-16.5 min followed by 2.5-minute hold. The mass spectrometer used for this study was Orbitrap Fusion Lumos Tribrid Mass Spectrometer (Thermo Fisher Scientific, Waltham, MA, USA) with heated electrospray ionization (HESI) source. Data were acquired in positive and negative-ion mode at a resolution of 120,000 within the mass range of 60– 850 *m/z*. The HESI source was operated at a voltage of 3000 V in negative-ion mode and 3500V in positive-ion mode. Source parameters such as sheath gas and auxiliary gas were set to 50 arbs and 15 arbs respectively. The ion transfer tube was heated to 330°C; the vaporizer temperature was set to 350°C. For metabolite confirmation, an in-house spectral library was used to verify mass detection at MS1 level and matching retention time ^69^.

#### Data analysis

MSI processing was carried out in LipostarMSI (Molecular Horizons, Italy; Tortorella et al.). Thermo RAW (.raw) files were imported and converted to imzML files using ProteoWizard (3.0.22317) as an intermediate within the in-built conversion tool. The imaging files were then imported to LipostarMSI using the following parameters: 5ppm m/z tolerance, peak intensity >1%, peak detection frequency >1% and minimum spatial chaos >0.7. Hotspot removal was applied using the 99% quantile and no denoising was applied. TIC normalisation was applied to the images.

Confirmatory identifications of reported metabolites were achieved by comparing serial bulk tissue analysis with a previously generated in-house spectral library ^69^. For the LC-MS analysis, Thermo XCalibur QualBrowser and Progenesis QI (v.3.0., Waters) was used for data analysis of mass spectra. In Progenesis QI, sample data and QC data was imported, and aligned with in-house library data for retention time and MS1 mass matching.

---

**Figure S1.**
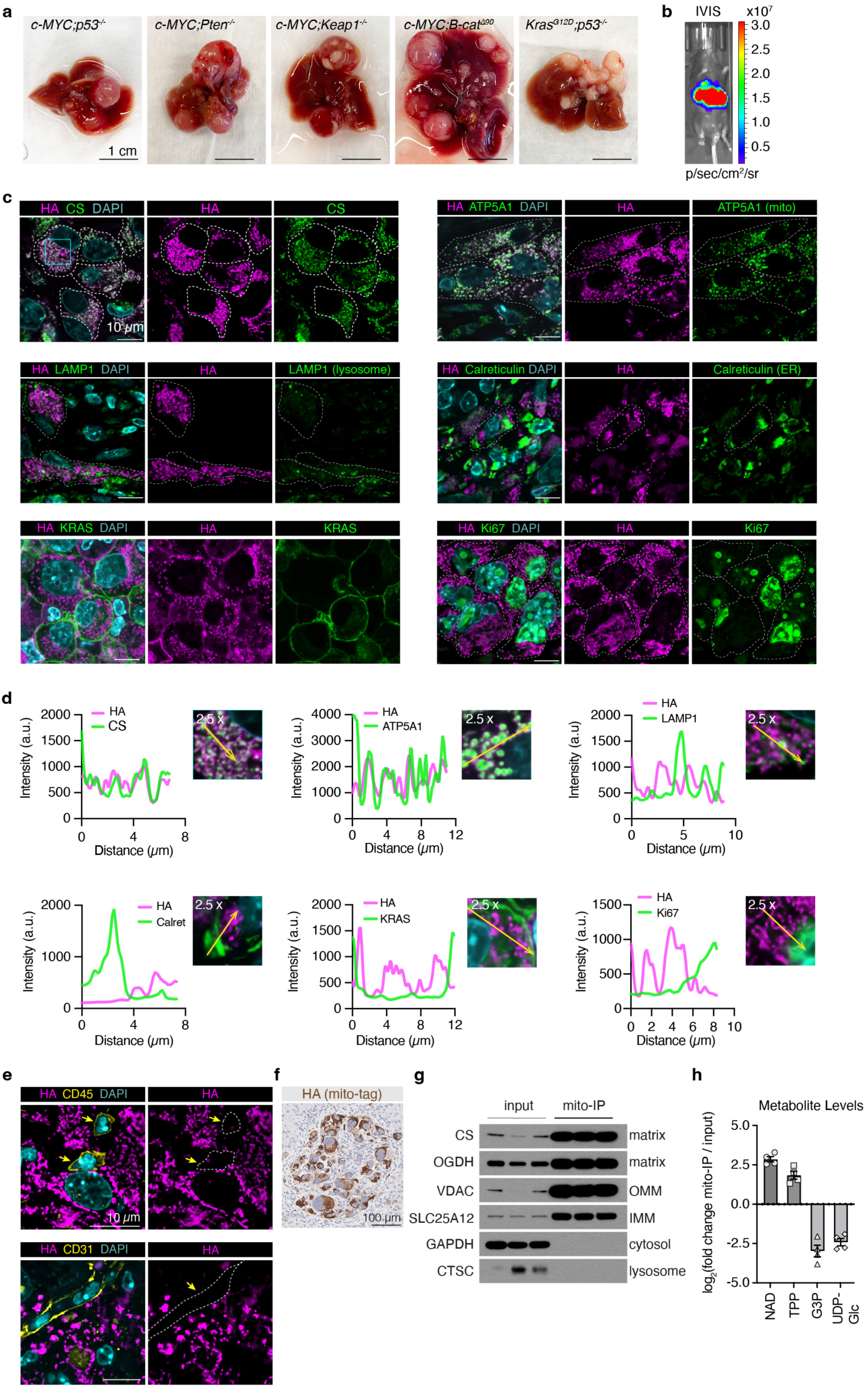
Mitochondrial localization of mito-tag in PLC tumors. a. Example images of genetically distinct PLC tumors generated via HDTVi method using mito-tag constructs. Scale bar is 1 cm. b. In vivo imaging software (IVIS) image of a live mouse with a liver tumor expressing mito-Luciferase tag. c. Immunofluorescence co-staining of mito-tag (HA-tag), DAPI (nuclear counterstain) and mitochondrial markers citrate synthase (CS) and ATP5A1; also other cellular compartment markers LAMP1 (lysosome), Calreticulin (ER), KRAS (plasma membrane localized proliferative marker) and Ki67 (nuclear localized proliferative marker). Scale bar is 10 *μ*m. d. Line scan plots showing co-localization of mito-tag (HA-tag) with mitochondrial marker CS and ATP5A1 as well as exclusion from other cellular compartments. 2.5X magnified views show line scan routes across the arrow. e. Immunofluorescence staining of mito-tag (HA-tag) and immune cell marker, CD45 (top panel) and endothelial cell marker CD31 (bottom panel). DAPI as nuclear counterstain f. Immunohistochemistry staining for mito-tag (anti-HA) shows tumor area containing mito-tag/HA^+^ cancer cells and unlabeled infiltrating cells, indicating cell type-specific labeling within the heterogenous tumor microenvironment. Scale bar is 100 *μ*m. g. Immunoblot showing enrichment of proteins localized to various sub-compartments of mitochondria, including mitochondrial matrix, inner mitochondrial membrane (IMM) and outer mitochondrial membrane (OMM). Of note, cytosolic (GAPDH) and lysosomal proteins (CTSC: cathepsin C) are depleted in mito-immunopurified fraction (mito-IP). h. Plot from metabolomics data displaying enrichment of mitochondrially concentrated metabolites nicotinamide dinucleotide (NAD) and thiamine pyrophosphate (TPP) in mito-IP pool, while other metabolites including Glycerol-3-phosphate (G3P) and Uridine diphosphate-glucose (UDP-Glc) are depleted, showing efficiency of mitochondrial enrichment.

**Figure S2.**
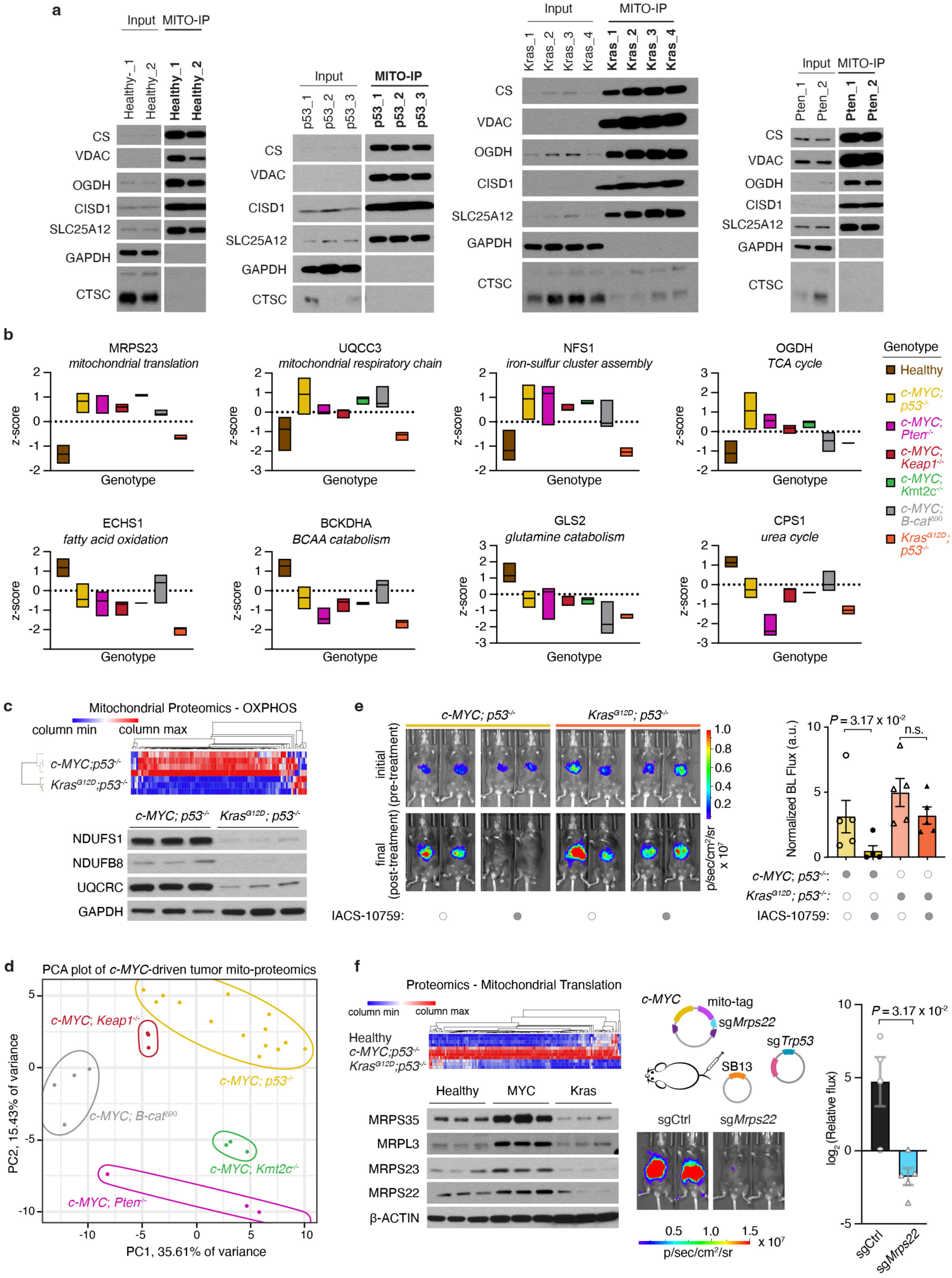
Mitochondrial translation and OXPHOS are metabolic vulnerabilities for *c-MYC; p53*^-/-^ driven PLC tumors. a. Example immunoblots showing enrichment of mitochondrial protein and depletion of other compartments (i.e., cytosol and lysosomes) from healthy liver and tumor mito-immunopurified fractions. p53: *c-MYC; p53 ^-/-^*, Pten: *c-MYC; Pten ^-/-^*, Kras: *Kras^G12D^; p53 ^-/-^*. Mitochondrial markers: CS (citrate synthase), VDAC, OGDH, CISD1 and SLC25A12. Cytosolic marker: GAPDH. Lysosomal marker: CTSC (cathepsin C). Input designates whole tumor lysate, MITO-IP is the mito-immunopurified fraction. b. Box plots for example proteins in mitochondrial translation (MRPS23), mitochondrial respiratory chain (UQCC3), iron-sulfur cluster assembly (NFS1), TCA cycle (OGDH), fatty acid oxidation (ECHS1), branched chain amino acid (BCAA) catabolism (BCKDHA), glutamine catabolism (GLS2), urea cycle (CPS1); and their expression changes as median normalized z-scores. Median is indicated as line across the box; minimum and maximum values define upper and lower boundaries c. Heatmap for OXPHOS proteins in *c-MYC; p53^-/-^* and *Kras^G12D^; p53^-/-^* models. Rows are clustered by hierarchical clustering. Column min and max are distributed across color scale (log scale). Immunoblot for select OXPHOS components in *c-MYC; p53^-/-^* and *Kras^G12D^; p53^-/-^* tumors (three independent tumors of each genotype) shows enrichment in *c-MYC*-driven tumors. GAPDH was used as control (non-OXPHOS protein). d. PCA plot for mitochondrial proteome of *c-MYC*-driven PLC models e. IACS-10759 treatment of *c-MYC; p53^-/-^* and *Kras^G12D^; p53^-/-^* tumor bearing mice. IVIS images of pre- and post-treatment tumor bearing mice (two representative mice per group) are displayed. Normalized bioluminescence (BL) flux data post-treatment is plotted as bar graph, showing *c-MYC*-specific tumor suppressive effect of IACS-10759. *P*-values are calculated by Mann-Whitney U-test with 95% confidence interval (CI). n.s., not significant. f. Heatmap for mitochondrial translation machinery components in *c-MYC; p53^-/-^* and *Kras^G12D^; p53^-/-^* models. Columns and rows are clustered by hierarchical clustering. Column min and max are distributed across color scale (log scale). Immunoblot for select mitochondrial translation components in healthy liver tissue, *c-MYC; p53^-/-^* and *Kras^G12D^; p53^-/-^* tumors is displayed on the left bottom panel. Middle top panel: Experimental design for generating *Mrps22*-depleted *c-MYC; p53^-/-^* tumors. IVIS images of representative mice bearing tumors are shown below. Plot on the right displays quantification for control (sgCtrl) and *Mrps22*-depleted (sg*Mrps22*) *c-MYC; p53^-/-^* PLC tumors. *P*-value is calculated by Mann-Whitney U-test with 95% confidence interval.

**Figure S3.**
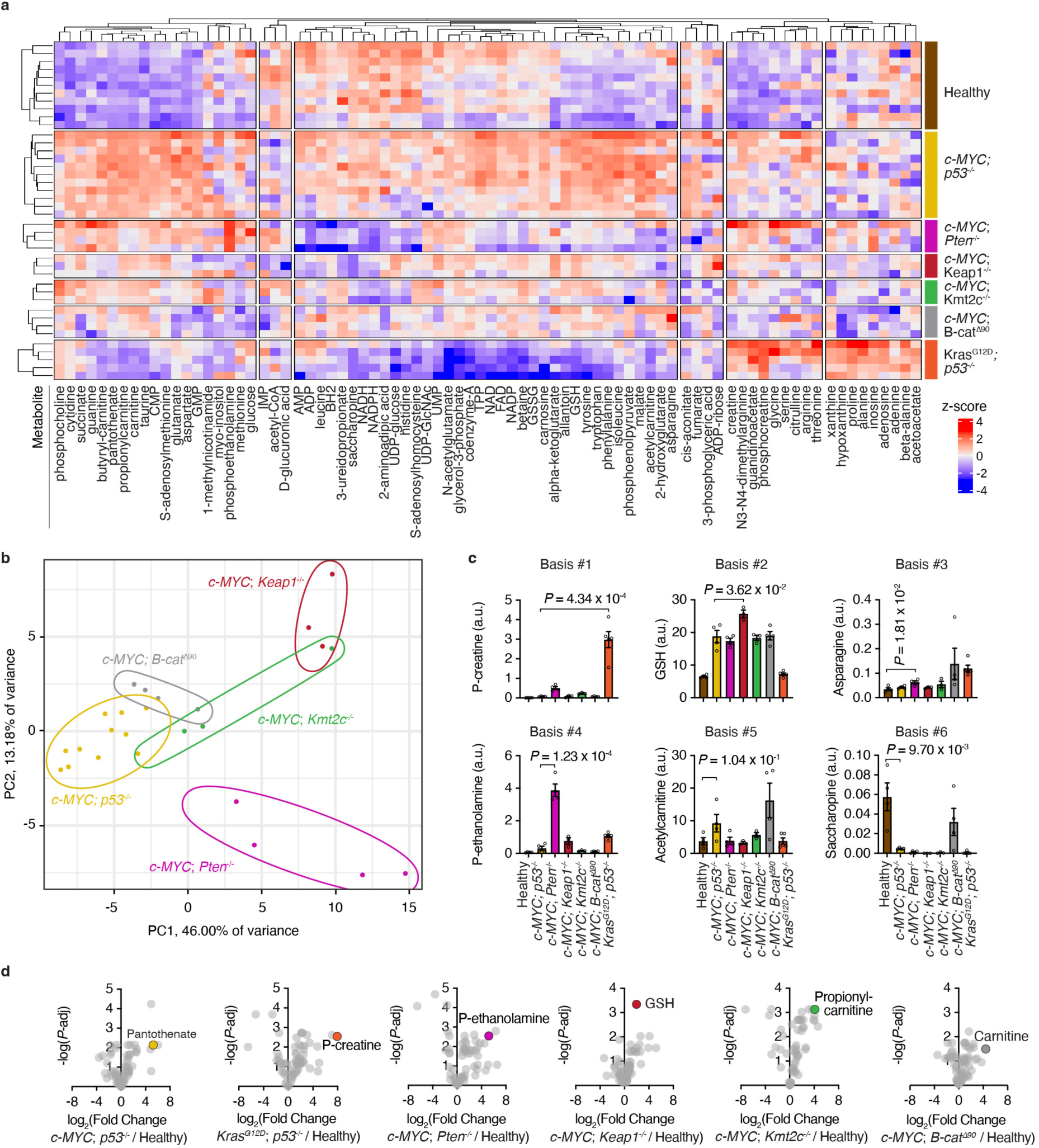
Mitochondrial metabolites altered in an oncogene- and tumor suppressor-specific manner. a. Heatmap showing median normalized changes in mitochondrial metabolites isolated from healthy livers and different tumor models, as indicated on the right by colored bars. Rows and columns are clustered by hierarchical clustering. b. PCA plot for mitochondrial metabolome of *c-MYC*-driven PLC tumor models c. Bar graphs showing levels of select metabolites enriched in each NMF basis. Student’s t-test for comparison with 95% confidence interval. GSH: glutathione-reduced. a.u.: arbitrary units. P-creatine: phosphocreatine; P-ethanolamine: phosphoethanolamine. d. Volcano plots for mitochondrial metabolite profiling, comparing tumors and healthy liver tissue (log_2_-scale). -log *P*-value was calculated by Student’s t-test and FDR-adjusted by Benjamini-Hochberg. Confidence interval is 95%.

**Figure S4.**
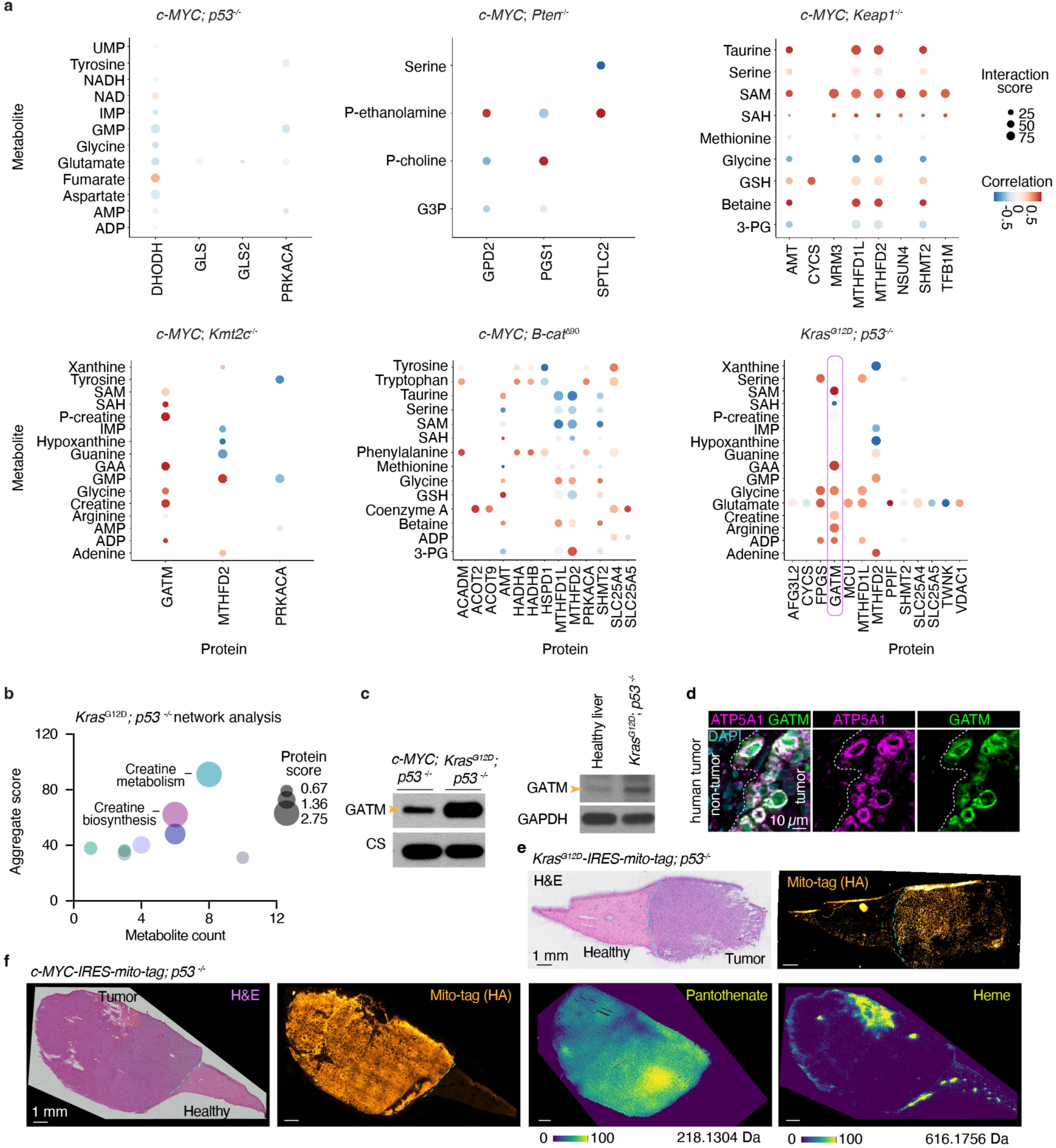
Creatine metabolism is a metabolic liability for *Kras*-mutated tumors. a. Dot plot of integrative network analysis displaying metabolites and proteins that are part of the same metabolic pathway. Protein-metabolite correlation values (color) and interaction scores (size) are indicated. GATM and its correlated metabolites in the creatine metabolism pathway are indicated in *Kras^G12D^; p53^-/-^* dataset. b. Integrative network analysis results of *Kras^G12D^; p53^-/-^* tumors showing enrichment of creatine metabolism. c. Immunoblot showing GATM expression in *c-MYC; p53^-/-^* and *Kras^G12D^; p53^-/-^* tumor lysates (left panel); right panel shows GATM immunoblot for lysates from healthy adjacent liver tissue and *Kras^G12D^; p53^-/-^* tumor. GAPDH as loading control. Arrow indicates the GATM band. d. Human ICC patient tissue, carrying *KRAS* mutation, stained for GATM and ATP5A1 (mitochondrial marker). Tumor boundary demarcated by dotted line. DAPI: nuclear counterstain. Scale bar is 10 *μ*m. e. H&E staining and mito-tag (HA-tag) immunofluorescence images of a serial section from liver tissue with *Kras^G12D^; p53^-/-^* tumor used for MALDI-MSI-imaging in Figure 2e. Tumor and healthy liver boundary is demarcated by blue dashed lines. Scale bar is 1 mm. f. H&E staining, mito-tag (HA-tag) immunofluorescence and MALDI-MSI images of a liver tissue with *c-MYC; p53^-/-^* tumor. Pantothenate [M+H^-^] and heme [M+H^+^] molecular images are displayed. Molecular ion mass is indicated below the image. Signal intensity is distributed across 0-100% color scale (color bar below). Da: Dalton. Scale bar is 1 mm.

**Figure S5.**
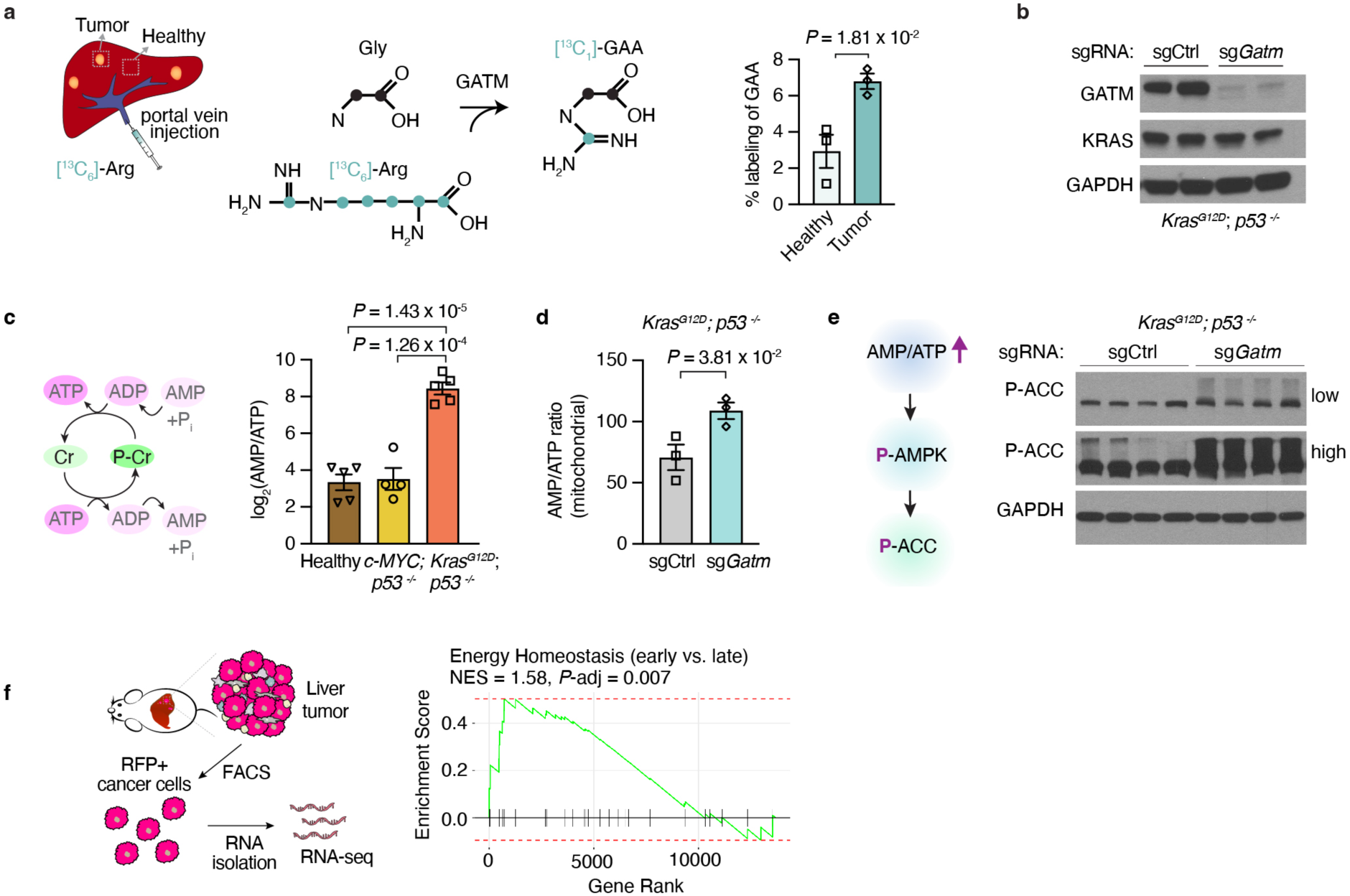
Energy homeostasis in *Kras*-mutated tumors. a. Experimental overview of isotope tracing by isotope labeled [^13^C_6_]-Arginine via portal vein injection. Isotope labelled [^13^C_6_]-guanidinoacetate (GAA) is formed by GATM activity. Per cent labeling of GAA in tumor and healthy adjacent tissue is plotted in the bar graph. *P*-value is calculated by Mann-Whitney U-test with 95% confidence interval. b. Immunoblot showing depletion of GATM in *Kras^G12D^;p53^-/-^* tumors (two representative samples from sgCtrl and sg*Gatm* group) by HDTVi-based method. KRAS and GAPDH were used as loading controls. c. Phosphocreatine (P-Cr)-Creatine (Cr) cycle to generate ATP. AMP/ATP ratios in log_2_ scale across healthy liver tissue, *c-MYC; p53^-/-^* and *Kras^G12D^; p53^-/-^* tumor samples d. Mitochondrial AMP/ATP ratios of *Kras^G12D^; p53^-/-^* control (sgCtrl) and *Gatm*-depleted (sg*Gatm*) tumors e. Schematic showing AMPK induction by increased AMP/ATP ratios and downstream ACC phosphorylation. On the right, phospho-ACC (P-ACC) immunoblot from *Kras^G12D^; p53^-/-^* control (sgCtrl) and *Gatm*-depleted (sg*Gatm*) tumor lysates (four independent tumors of each genotype). Low and high exposure of the blot are provided for P-ACC. GAPDH is displayed as loading control. f. Strategy for isolation of RFP^+^ cancer cells from *Kras^G12D^; p53^-/-^* tumors by fluorescence-associated cell sorting (FACS), RNA isolation and RNA-seq. Right panel: gene set enrichment analysis of *Kras^G12D^; p53^-/-^* tumors showing enrichment of ‘Energy Homeostasis’ in early-stage tumors as compared to late-stage tumors

**Figure S6.**
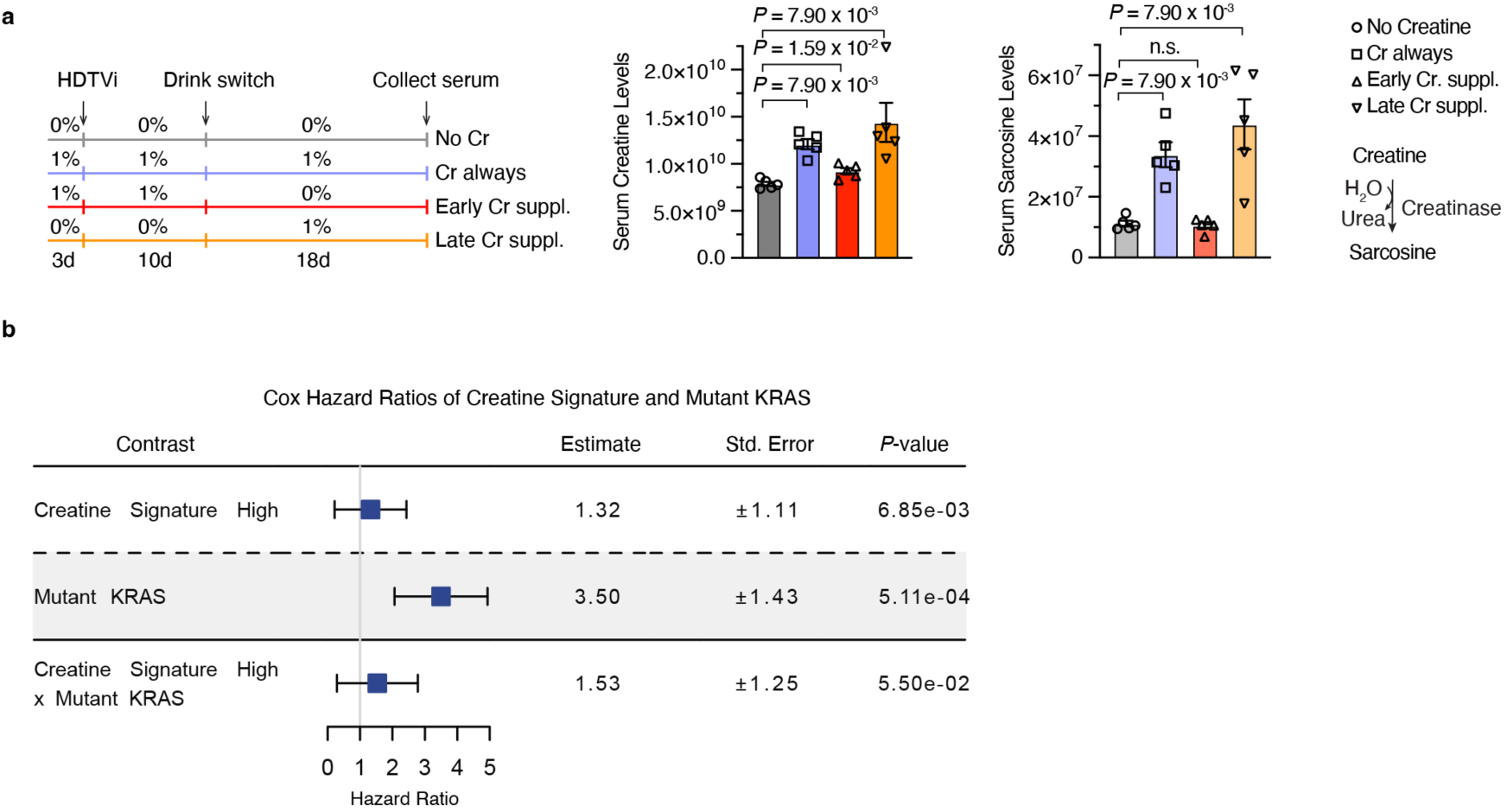
Creatine supplementation and Cox hazard ratios for creatine signature in the patient cohort. a. Overview of creatine supplementation and serum collection for metabolite profiling by LC/MS. Bar graphs show levels of serum creatine and sarcosine, a creatine degradation product. Data are mean± SEM, comparison by Student’s t-test with 95% CI. b. Table for Cox hazard ratios of creatine synthesis and mutant *KRAS* status in ICC patients. Synergistic effect of high creatine signature and mutant *KRAS* is noted in this analysis.

**Figure S7.**
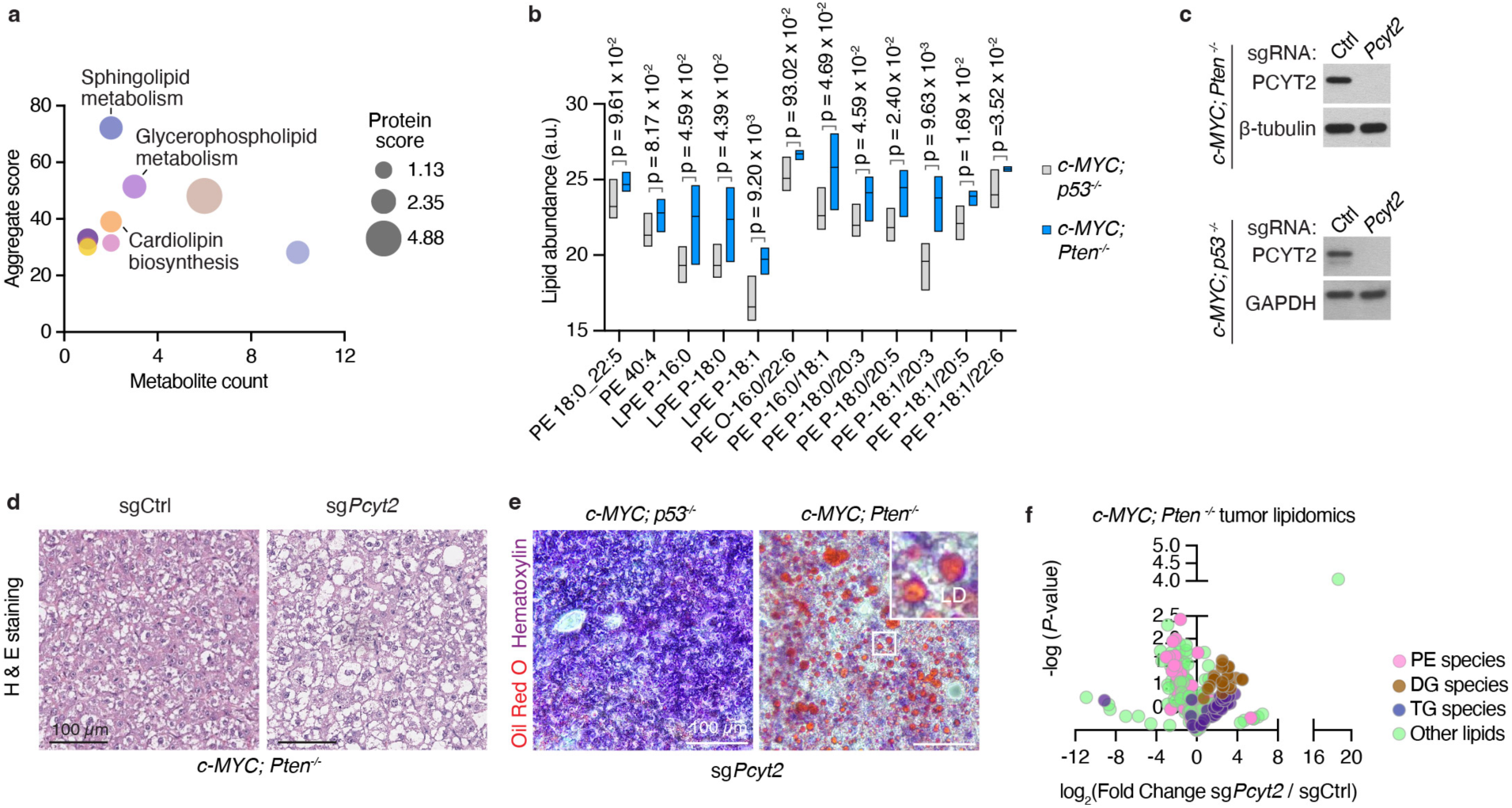
Lipid metabolism-related phenotypes *in c-MYC; Pten^-/-^* PLC tumors. a. Integrative network analysis for *c-MYC; Pten^-/-^* tumors showing enrichment of lipid metabolic processes in *c-MYC; Pten^-/-^* tumors b. Box plot for lipidomics data comparing *c-MYC; Pten^-/-^* and *c-MYC; p53^-/-^* tumors. Median is indicated as line across the bar; minimum and maximum values define upper and lower boundaries. P-values are calculated by Student’s t-test; confidence interval 95%. c. Immunoblot shows depleted PCYT2 levels in *c-MYC; Pten^-/-^* tumors. β-tubulin as loading control. Lower immunoblot panels shows PCYT2 depletion in *c-MYC; p53^-/-^* tumor background; GAPDH as loading control. d. H&E staining for *c-MYC; Pten^-/-^* tumors, control (sgCtrl) and *Pcyt2*-deficient (sg*Pcyt2*). Scale bar is 100 *μ*m. e. Oil red O staining for *Pcyt2*-depleted *c-MYC; p53^-/-^* and *c-MYC; Pten^-/-^* tumors. Scale bar is 100 *μ*m. f. Volcano plot for lipidomic analysis of *c-MYC; Pten^-/-^* tumors comparing *Pcyt2*-deficient and control tumors. PE: phosphotidylethanolamine, DG: diacylglyceride, TG: triacylglyceride

**Figure S8.**
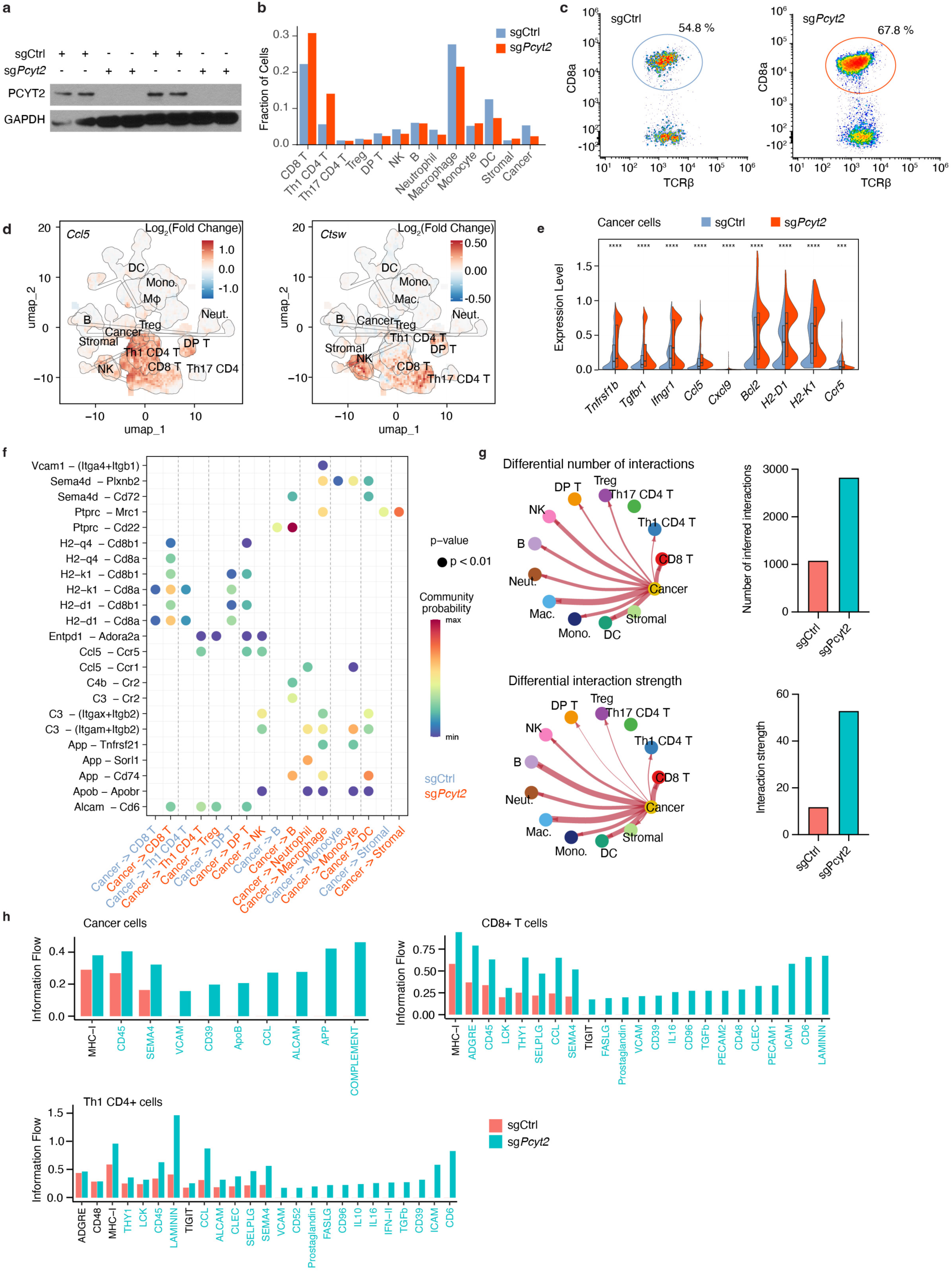
Activation markers are upregulated in immune cells infiltrating *Pcyt2*-deficient *c-MYC; Pten^-/-^* PLC tumors. a. Immunoblots for *c-MYC; Pten^-/-^* tumors included in the sc-RNA seq experiment. Four tumors of each sgCtrl and sg*Pcyt2* genotype. GAPDH used as loading control. b. Fraction of cell types detected in *Pcyt2*-depleted (sg*Pcyt2*) or control (sgCtrl) tumors. c. Flow cytometry analysis of TCRβ+ CD8a+ T-cells in control (sgCtrl) and *Pcyt2*-depleted (sg*Pcyt2*) tumors d. UMAP plots showing log_2_ fold change in expression (sg*Pcyt2* vs. sgCtrl) of immune cell activation (*Ccl5* and *Ctsw*) e. Differential gene expression analysis of immune signaling receptors in cancer cells f. Dot plot showing the results of CellChat algorithm identifying immune signaling interactions between cancer and infiltrating immune cells. Presence of a dot indicates significance at p<0.01, colors represent community probability g. Differential analysis of number and strength of intercellular interactions by CellChat. Line thickness represents the number or strength of interaction, accordingly. Bar plots show the numeric data from these analyses. h. Information flow through cell surface molecules in cancer cells, CD8^+^ and Th1 CD4^+^ T cells. Data from sgCtrl and sg*Pcyt2 c-MYC; Pten^-/-^* tumors.

**Figure S9.**
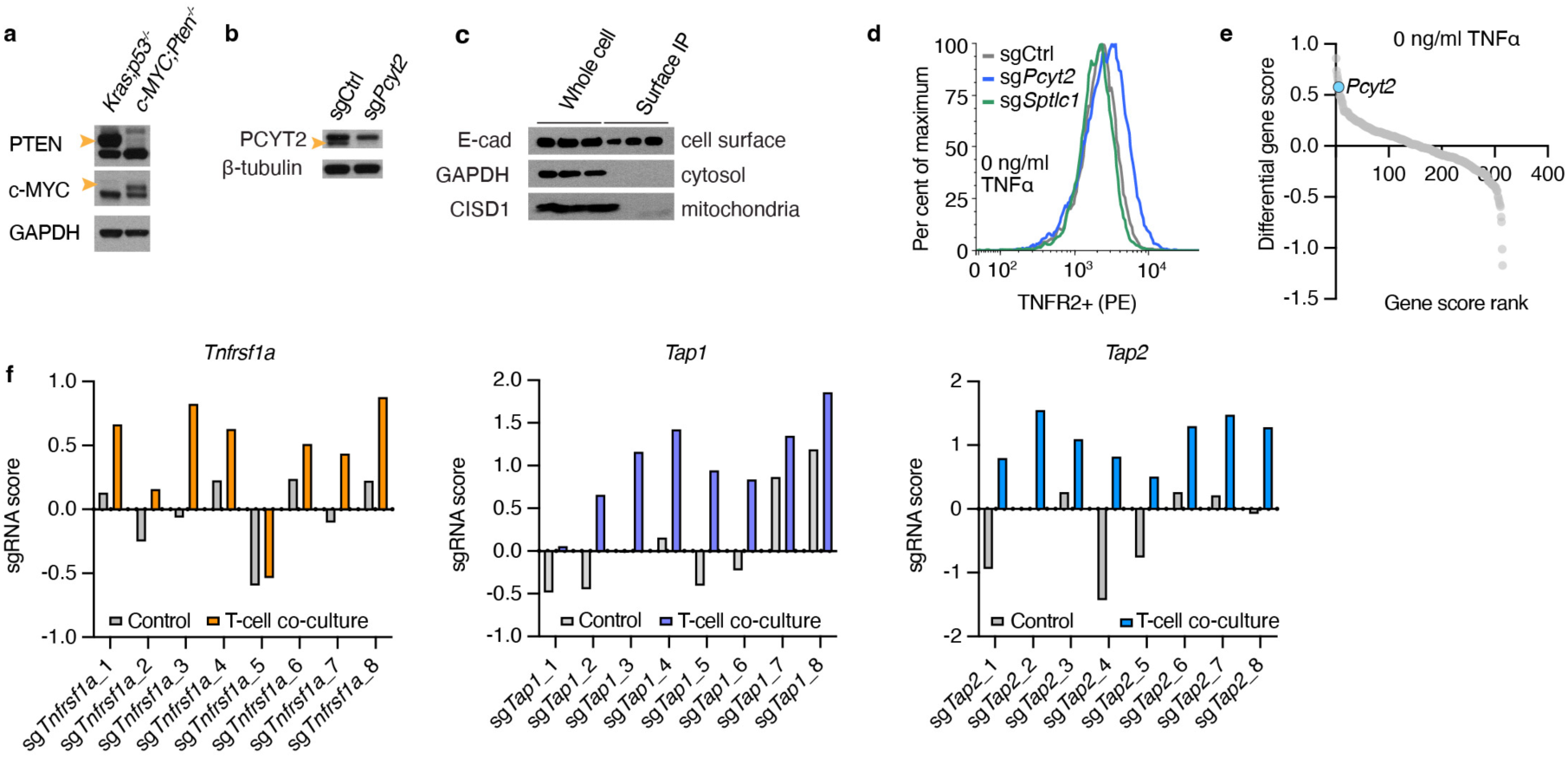
*Pcyt2* loss affects surface TNFR2 levels. a. Immunoblot verifying *c-MYC; Pten^-/-^* liver cancer cell line. HY15549 *Kras*; *p53^-/-^* cell line was used as control. GAPDH for loading control. b. Immunoblots for PCYT2-depleted cells used for cell surface proteomics. c. Immunoblot demonstrates presence of E-cadherin (E-cad), and depletion of cytosolic GAPDH and mitochondrial CISD1 in cell surface immunoprecipitated fraction. d. Flow cytometry analysis of baseline TNFR2 surface expression in untreated sgControl, sg*Pcyt2* and sg*Spltc1* (used as a control for depletion of another membrane lipid class) *c-MYC; Pten^-/-^*. PE: phycoerythrin. e. FACS-based screen to identify lipid metabolism genes that affect the stability of TNFR2 in the plasma membrane of *c-MYC; Pten^-/-^* cells (untreated baseline). Live and transduced cells were incubated with phycoerythrin tagged antibody against TNFR2. High- and low-fluorescence populations were collected and amount of sgRNA was compared. Ranks of gene scores between TNFR2^hi^ and unsorted control population is plotted. *Pcyt2* was amongst the top 10 genes. f. sgRNA scores of *Tnfrsf1a*-, *Tap1*- and *Tap2*-targeting targeting sgRNAs in control and OT-I CD8^+^ T-cell co-culture CRISPR screens.

**Table S1.** Mitochondrial proteomics data. Median normalized z-scores are listed.

**Table S2.** Metabolic network analysis results.

